# Finite element model predicts micromotion-induced strain profiles that correlate with the functional performance of Utah arrays in humans and non-human primates

**DOI:** 10.1101/2025.04.10.648248

**Authors:** Adam M. Forrest, Nicolas G. Kunigk, Jennifer L. Collinger, Robert A. Gaunt, Xing Chen, Jonathan P. Vande Geest, Takashi D.Y. Kozai

## Abstract

**Objective:** Utah arrays are widely used in both humans and non-human primates (NHPs) for intracortical brain-computer interfaces (BCIs), primarily for detecting electrical signals from cortical tissue to decode motor commands. Recently, these arrays have also been applied to deliver electrical stimulation aimed at restoring sensory functions. A key challenge limiting their longevity is the micromotion between the array and cortical tissue, which may induce mechanical strain in surrounding tissue and contribute to performance decline. This strain, due to mechanical mismatch, can exacerbate glial scarring around the implant, reducing the efficacy of Utah arrays in recording neuronal activity and delivering electrical stimulation.

**Approach:** To investigate this, we employed a finite element model (FEM) to predict tissue strains resulting from micromotion.

**Main Results:** Our findings indicated that strain profiles around edge and corner electrodes were greater than those around interior shanks, affecting both maximum and average strains within 50 µm of the electrode tip. We then correlated these predicted tissue strains with *in-vivo* electrode performance metrics. We found negative correlations between 1 kHz impedance and tissue strains in human motor arrays and NHP area V4 arrays at 1-mo, 1-yr, and 2-yrs post-implantation. In human motor arrays, the peak-to-peak waveform voltage (PTPV) and signal-to-noise ratio (SNR) of spontaneous activity were also negatively correlated with strain. Conversely, we observed a positive correlation between the evoked SNR of multi-unit activity and strain in NHP area V4 arrays.

**Significance:** This study establishes a spatial dependence of electrode performance in Utah arrays that correlates with tissue strain.

## 1. Introduction

Brain-computer interfaces (BCIs) are advanced technologies that can be integrated into therapeutic strategies to restore motor and sensory function in individuals with central nervous system injuries and disorders [1, 2]. BCIs have previously employed arrays of ∼100 intracortical microelectrode shanks implanted in the motor cortex to capture neural signals by recording action potentials from surrounding neurons, which can then be decoded to control devices, such as robotic limbs [3–7]. Additionally, in the somatosensory and visual cortices, microelectrode arrays have been used to deliver electrical stimulation and restore artificial tactile and visual sensations [8–13]. A critical consideration for implanted BCIs is the longevity of these devices [14–21]. While intracortical arrays can record action potentials for years, their ability to record from nearby neurons is highly variable [22, 23] and diminishes over time [15, 19, 24–27]. Although researchers have investigated new electrode shank designs to improve longevity [28–38], the influence of neighboring shanks on each other remains underexplored.

Designs of microelectrode arrays influence the degree of tissue response following microelectrode implantation, in turn impacting device longevity. Acute tissue damage during implantation disrupts the blood-brain barrier, reducing nutrient delivery to the disrupted tissue, releasing plasma proteins into the parenchyma, and triggering an inflammatory response that leads to an electrically insulating glial scar around the microelectrodes [39–43]. Activated microglia and astrocytes at the glial scar release cytokines that promote a neurodegenerative environment at the electrode-tissue interface further reducing neuronal activity and signal detectability [44–52]. This neuroinflammation is exacerbated by strain concentrations in the tissue caused by micromotions of the electrode relative to the brain [53–55]. Behavioral head movements, respiration, and blood vessel pulsations contribute to brain movement [56–59]. Because typical electrodes are 8 orders of magnitude stiffer than brain tissue, this relative motion generates mechanical strain in the surrounding parenchyma [60], which increases scarring around implants and impairs BCI performance [61–68].

Mechanical strain can activate neurons and glial cells via mechanosensitive ion channels [53, 54, 69]. In microglia, these channels can direct migration towards stiffer substrates and prompt phagocytic activity of cellular debris [58, 59, 70, 71]. Similarly, astrocytic inflammatory responses are mediated through mechanosensitive ion channels [43, 72, 73]. Reducing mechanical strain in the surrounding tissue could minimize glial activation and scarring, thereby promoting an improved electrode-tissue interface [65, 66, 74, 75]. Neurons can be directly damaged by mechanical injury during implantation, which is evidenced by altered cellular morphology and elevated calcium levels, potentially impacting the ability to reliably record neuronal activity [65, 76]. Damage to blood vessels may also reduce metabolic support to nearby neurons and contribute to local neuronal dysfunction, hampering signal acquisition and BCI performance [77–79]. As the number of shanks increases [37, 80–82], the cumulative mechanical strain within the tissue can become more pronounced [34, 83], exacerbating these effects by further compromising vascular integrity, increasing the extent of glial scarring, and reducing the overall functionality and longevity of microelectrodes. This heightened strain may also lead to greater variability in signal acquisition across the array, complicating the interpretation of neural signals and the effectiveness of the BCI system.

Finite element models (FEMs) have shown that linear electrodes, characterized by a single shank with multiple electrode sites along its length, exhibit high strain profiles near the electrode tip due to micromotions [84–90]. This strain concentration has been linked to neuroinflammation and the foreign-body response, limiting the longevity and reliability of these devices. However, strain profiles around bed-of-needle planar-style Utah arrays, characterized by rectangular or square base substrates with equidistantly spaced electrodes, have not been thoroughly characterized. Shank-to-shank distance alters tissue strain profiles during insertion [34], but the impact of micromotions on strain distributions in Utah arrays remains unclear. Understanding how micromotion-induced tissue strains affect device longevity is crucial for designing long-lasting multi-shank microelectrode arrays.

This study investigates how Utah array geometry influences tissue strain and recording performance. Using a FEM, micromotion-derived strain was predicted throughout the surrounding tissue and then correlated with device performance metrics, including impedance, peak-to-peak waveform voltage, and signal-to-noise ratio (SNR) in both humans and non-human primates (NHPs). The distinct geometry of Utah arrays suggests that tissue strain profiles will vary based on electrode location within the array. For example, edge and corner electrodes may experience different mechanical interactions with surrounding brain tissue compared to those in the interior. The array’s geometry could also influence vascular and ischemic injuries, which are known to impact device performance. Vascular injury may be more pronounced at the edges, where blood vessels are more susceptible to implantation damage due to tissue dimpling [37, 91]. Conversely, ischemic injury could be greater at the array’s center, where the distance to undamaged blood supply is greater. While changes in performance metrics over the implant’s lifetime have been documented, the spatial dependence of these measures within Utah arrays remains underexplored. Addressing this gap will enhance understanding of how electrode positioning within the array impacts overall device performance, longevity, and the health of surrounding tissue.

## 2. Methods

### 2.1 Finite Element Modeling

#### 2.1.1 Geometry

Electrode array geometries were designed to replicate the dimensions of the Utah arrays used in the electrophysiological recordings (Blackrock Neurotech). Three array geometries were modeled: 100- electode arrays in a 10x10 grid, 60-electrode arrays in a 10x6 grid, and 64 electrode arrays in an 8x8 grid. Each grid of electrodes had the same 400 µm spacing. 10x10 arrays were modeled with a square 4mm x 4mm x 0.2mm base substrate. Electrode shanks were defined by tapering from a 150 µm × 150 µm square at the base to a 10-µm diameter circle at 1.49 mm from the base. To form the tip, a quarter ellipse with a major radius of 10 µm and a minor radius of 5 µm was revolved, resulting in a total shank length of 1.5 mm. 10x6 and 8x8 arrays were modeled in the same fashion, but utilized a rectangular 4mm x 2.4mm x 0.2mm base substrate and a square 3.2mm x 3.2mm x 0.2mm base substrate, respectively. Surrounding cortical brain tissue was modeled as a rectangular prism extending 1 mm beyond the electrode tips and 2 mm outward from the base on all sides. This tissue was modeled as a separate entity in fixed contact with the electrode shanks and substrate (Fig. 1a).

**Figure 1.**
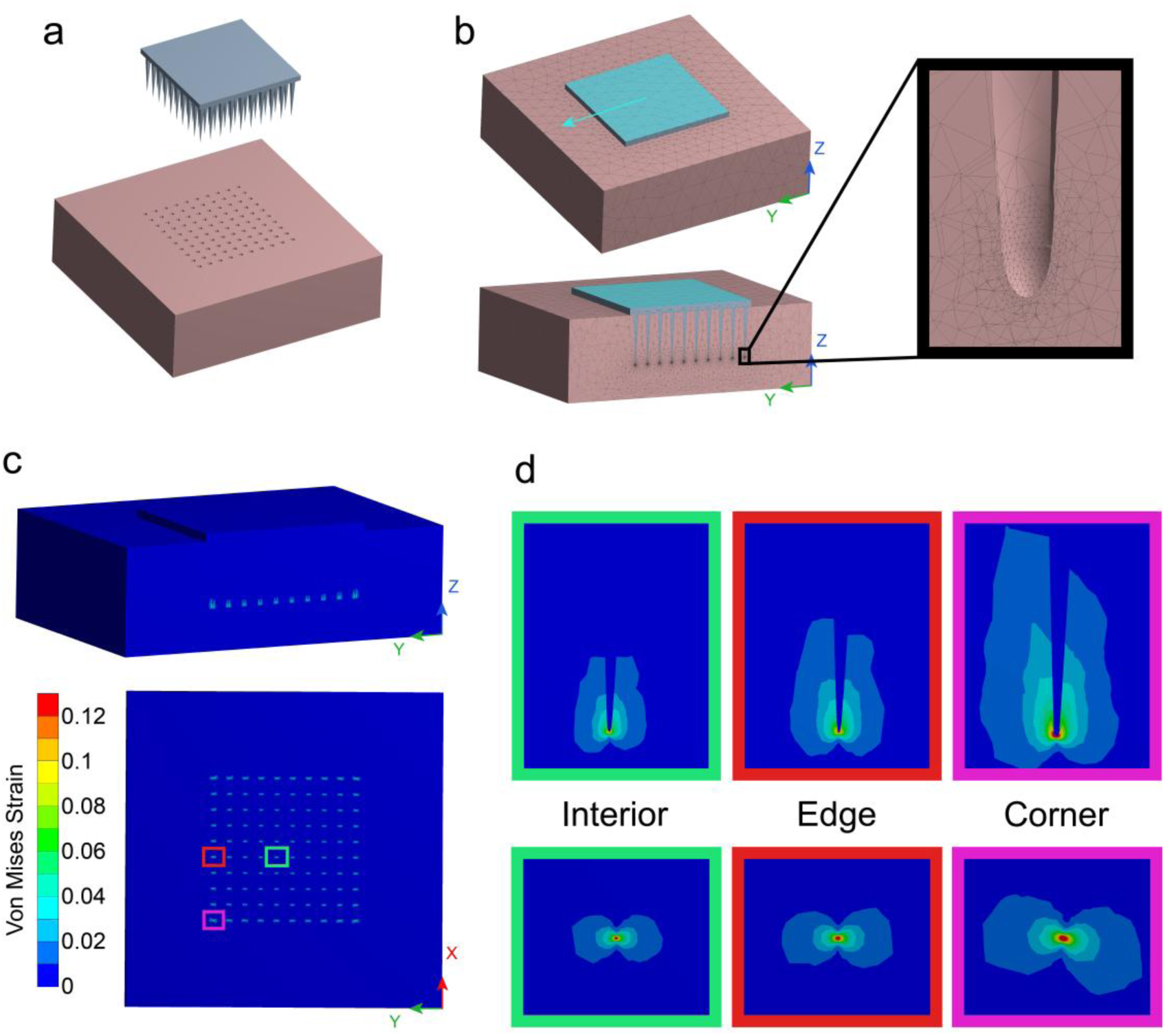
Finite element model predicts strains in brain tissue surrounding implanted Utah array that result from micromotions. (a) Geometry of 10x10 array (top) and brain tissue (bottom). (b) Mesh of 10x10 array embedded in cortical tissue with applied boundary conditions. The top face of the array was prescribed a displacement of 10µm in the Y direction. The bottom face of the brain tissue was held fixed. Sliced model (bottom) shows the fine mesh surrounding the tips of the shanks, further shown in inset (right). (c) Contour plots of von Mises strain. Model was sliced in YZ plane (top) and XY plane (bottom) to uncover the tips of the electrodes (d) Contour plots surrounding an example interior, edge, and corner electrode. Color bar is same as in (c).

Additional geometries were generated with the electrodes terminating to an exact point. These models were untenable as the finite element models would not reach mesh convergence due to infinite point stresses at the shank tip.

#### 2.1.2 Material Properties

The electrode array was assumed to be silicon with linear isotropic properties. Based on previous studies [85, 86], we used a Young’s modulus of 200 GPa and a Poisson’s ratio of 0.278. Material properties of the brain were assumed to be nonlinear and were modeled using a 1^st^ order Ogden hyperelastic model (Equation 1) with the strain-energy equation,

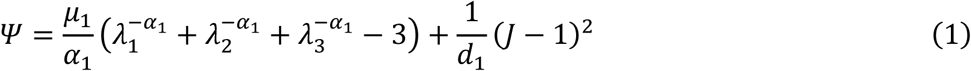

where 𝜆_𝑖_ are the principal stretches, 𝐽 is the determinant of the elastic deformation gradient, and 𝜇_1_, 𝛼_1_, and 𝑑_1_ are the material properties. Budday et. al. previously characterized material properties of human cortical tissue using this model [92]. Namely, 𝜇_1_ = 150.5 𝑃𝑎, 𝛼_1_ = 19, and 𝑑_1_ = 6.65 × 10^−5^ 𝑃𝑎^−1^.

#### 2.1.3 Boundary Conditions

The bottom face of the brain tissue, closest to the electrode tips, was spatially fixed (displacement of zero). To mimic the relative motion of the electrode with the cortical tissue, the top face of the electrode array (opposite the brain tissue) was prescribed a displacement of 10µm in the Y direction (Fig. 1b).

#### 2.1.4 Meshing

Meshes and finite element solutions were computed with ANSYS Mechanical 18.2 using a static structural problem type. Both the array and brain were meshed using 2^nd^ order tetrahedrons. Spheres of influence at the tip of each electrode were generated to refine the mesh near the shank tip. The sphere of influence was set to a radius of 10µm and constrained the mesh in this volume to have a maximal edge length of 1µm. This parameter choice was validated through a mesh convergence study. The full model for the 10x10 array and brain had a median element volume of 1.96 µm^3^ (IQR = 0.26 µm^3^ – 1014 µm^3^).

#### 2.1.5 Simulations and Strain Analysis

For each array geometry and applied boundary condition, von Mises strain was computed at each node. Von Mises strain is a scalar value that provides a single measure that incorporates axial and shear strains. Exported nodal displacements, strains, and locations were analyzed using MATLAB 2022a. Our region of interest consisted of all nodes within 50µm of the electrode tips, representing the approximate volume of cortical tissue where single units can be recorded [93, 94]. Maximum strain was defined as the maximal nodal strain within this ROI. To obtain a more representative metric over the entire ROI, nodal strains were averaged over 2.5µm bins extending out from the electrode tip. Binned strains were then normalized by the volume they occupied to compute the average strain in the 50µm sphere. Electrodes were grouped based on the number of rows away from the edge of the array, generating a series of concentric electrode “rings”. Corner electrodes were separated into their own group. Electrodes were also analyzed based upon their distance from the center of the array.

#### 2.1.6 Mesh Convergence

Mesh convergence was conducted to ensure our finite element solutions were mesh-independent. To improve computational efficiency, the mesh convergence study was carried out using a simplified array geometry with only one electrode shank. Meshes were refined using a 10µm sphere of influence surrounding the electrode tip with maximal element edge sizes of 3µm, 2µm, 1µm, 0.5µm, 0.3µm, and 0.2µm. Material properties and boundary conditions described previously were used. Average von Mises strain surrounding the electrode was then computed for each mesh. Maximum element edge size of 1µm was chosen as a tradeoff between accuracy and computational load, with less than 5% error compared to the finest mesh tested (Supplementary Figure 1a-b).

### 2.2 Neural Data Processing

#### 2.2.1 Human motor and somatosensory arrays

Data collected from Utah arrays implanted in clinical study participants (N=2 participants, n=4 motor arrays, n=4 somatosensory arrays; somatosensory arrays were used for microstimulation) was previously published by Flesher et. al. and others [9, 19, 25, 95]. For P2, 10x10 motor arrays were platinum and 10x6 somatosensory arrays were SIROF (sputtered iridium oxide film) coated. For P3, both 10x10 motor arrays and 10x6 somatosensory arrays were SIROF coated. Data were collected as part of an ongoing clinical trial of a sensorimotor brain-computer interface (NCT01894802) and conducted under an FDA Investigational Device Exemption with approval from the University of Pittsburgh Institutional Review Board. Methods for surgical implantation have been described previously [96–98]. Due to hardware limitations, recordings were not collected on all electrodes. For P2 10x10 motor arrays, three electrodes in each corner were unconnected. For P3 10x10 motor arrays, the four corner electrodes were unconnected. For all 10x6 somatosensory electrodes, connected electrodes form a checkerboard pattern. Impedance measurements at 1kHz were taken using the NeuroPort signal processor (Blackrock Neurotech). SIROF-coated electrodes with impedances above 1 MΩ and platinum electrodes with impedances above 2 MΩ were deemed nonfunctional and excluded from subsequent analysis.

Spontaneous electrophysiological recordings were taken at 30 kHz during a restful period where the participants relaxed in a quiet room. Extraction of waveforms were similar to previous methods by Hughes et al [19]. Electrophysiological recordings were filtered using a 2^nd^ order bandpass Butterworth filter with cutoff frequencies of 300Hz and 5000Hz. Recorded action potentials were detected using a threshold of 4.5 times the root-mean-square below the mean. Waveform snippets were generated by extracting a window 0.5ms before and 1ms after threshold crossing. The peak-to-peak voltage (PTPV) of each snippet was calculated by subtracting the minimum voltage from the maximum voltage. For each active electrode, an average was taken of the largest 2% of snippets to obtain a representative PTPV value. Only the largest snippets were used in order to estimate the maximum PTPV voltage that could be achieved. Signal-to-noise ratio (SNR) was computed by dividing the PTPV for that electrode by the noise floor, which was computed as twice the standard deviation of the recording after removing the snippets. Five successive sessions were analyzed for each time period of interest (1-mo, 1-yr, and 2-yrs post-implantation). For each array, impedance, PTPV, and SNR measures were first averaged across the five sessions before normalizing within the individual array (Supplementary Fig. 2-3). Session-to-session variability was assessed by analyzing the electrode standard deviation across the five successive sessions. We observed that edge electrodes had reduced session-to-session variability compared to interior electrodes for impedance and PTPV measures at 1-yr post-implantation (Supplementary Fig. 4).

We also assessed the similarity in bandpass-filtered recordings between neighboring electrodes. For each electrode, its bandpass-filtered recording was correlated with the bandpass-filtered recording on all neighboring electrodes (3-8 electrodes depending on location). The average of these 3-8 neighboring electrode correlations was then computed. These values were averaged within an array across five successive recording sessions for each time period. The average correlation values could theoretically vary between -1 and 1. However, due to the large number of positive correlations present, all averaged correlations were greater than zero.

#### 2.2.2 Non-human primate area V4 arrays

A second dataset from Utah arrays implanted in NHPs was also analyzed (N=2 animals, n=4 area V4 arrays) [27, 99]. For both animals, 8x8 V4 arrays were iridium-oxide coated. Methods for surgical implantation and data collection have been described previously. Impedance measurements at 1kHz were taken using the Impedance Tester in the Research Central Software Suite (Blackrock Neurotech). Visually evoked electrophysiological recordings were taken during a visual stimulation detection task. Snippet extraction and PTPV calculations were conducted in the same manner as with the human array data [19, 27]. Envelope multi-unit activity (eMUA) was also extracted as previously described [100]. Evoked SNR was computed by subtracting the average eMUA from baseline spontaneous activity from the peak trial-averaged eMUA during stimulation, and then dividing by the standard deviation of the baseline eMUA. Impedance, PTPV, and SNR measures were normalized within each individual array (Supplementary Fig. 5). Due to limited number of recording sessions, session-to-session variability was not assessed.

### 2.3 Statistics

Statistical analyses were performed in MATLAB2022a. Means across groups from the modelled data were compared using one-way ANOVA with Tukey’s HSD post-hoc testing. For electrophysiological data, linear regression and ANOVA assumptions of normality and homoscedasticity were not met across electrode location groups. Therefore, Kruskal-Wallis tests with post-hoc Dunn’s tests were used to determine whether a significant difference in medians across multiple groups exists. Similarly, linear correlation assumptions were violated when assessing the relationship between electrophysiological measures and modelled strain. As a result, Spearman’s rank-order correlation was conducted to test for significant correlation. For significant correlations (p<0.05) Spearman’s correlation coefficient (ρ) values are reported as a measure of the strength of the relationship. These values range from -1 to +1.

## 3. Results

We sought to determine the relationship between Utah array geometry, tissue strain, and the recording capabilities of these devices. We used FEMs to simulate micromotion-induced strain in brain tissue surrounding Utah arrays of various geometries, including 10x10, 10x6, and 8x8 grid configurations, to replicate those used in electrophysiological recordings. We then correlated these strain distributions with electrophysiological performance metrics—impedance, peak-to-peak waveform voltage, and SNR—collected from Utah arrays implanted in both humans and non-human primates.

### 3.1 Finite element model predicts micromotion-derived tissue strains at the edge of Utah arrays are greater than strains in the interior

We first asked how tissue strain varies from electrode to electrode across the Utah array. To this end, we used a finite element model to predict the tissue strains resulting from micromotions in the brain (Fig. 1a-b). Contour plots of the resulting von Mises strain for the 10x10 array are shown in Fig. 1c. We qualitatively observed increased tissue strains near the corner electrodes compared to the edge electrodes, both of which were greater than near interior electrodes (Fig. 1d). A mesh convergence study confirmed that the solution was independent of the mesh size (Supplementary Fig. 1a-b).

A sphere with radius 50µm centered at each electrode tip was defined as our region of interest (Fig. 2a). To obtain a strain metric that accounts for the entire ROI without bias from mesh density variations, nodal strains were averaged over 2.5µm bins extending out from the electrode tip (Fig. 2b). These binned strains were then normalized by the volume they occupied to calculate the average strain within the 50µm sphere. Distinct separation between the strain curves for corner, edge, and interior electrodes was observed. When assessing the average strain for individual electrodes, tissue near corner electrodes showed significantly higher strains compared to other edge electrodes, and edge electrodes had greater strains than interior electrodes (p<0.0001, one-way ANOVA with Tukey’s post-hoc, Fig. 2c-d). Additionally, the 2^nd^ and 3^rd^ outermost rings in the 10x10 array exhibited greater strains than the other interior electrodes. Further analysis revealed that grouping electrodes by concentric rings better distinguished strain differences than using distance from the array center (Fig. 2e). Similar trends were observed when analyzing the maximum strain within the ROIs, with tissue surrounding corner and edge electrodes showing significantly higher maximal strains compared to interior regions (p<0.0001, one-way ANOVA with Tukey’s post-hoc, Supplementary Fig. 6). Consistent results in the average strains surrounding 10x6 and 8x8 Utah arrays indicate that the spatial patterns were a consequence of the planar geometry of the array and not the number of electrodes (Supplementary Fig. 7a-b). These findings highlight the importance of electrode placement and array design in influencing tissue strain profiles, which can potentially impact overall device performance and tissue health.

**Figure 2.**
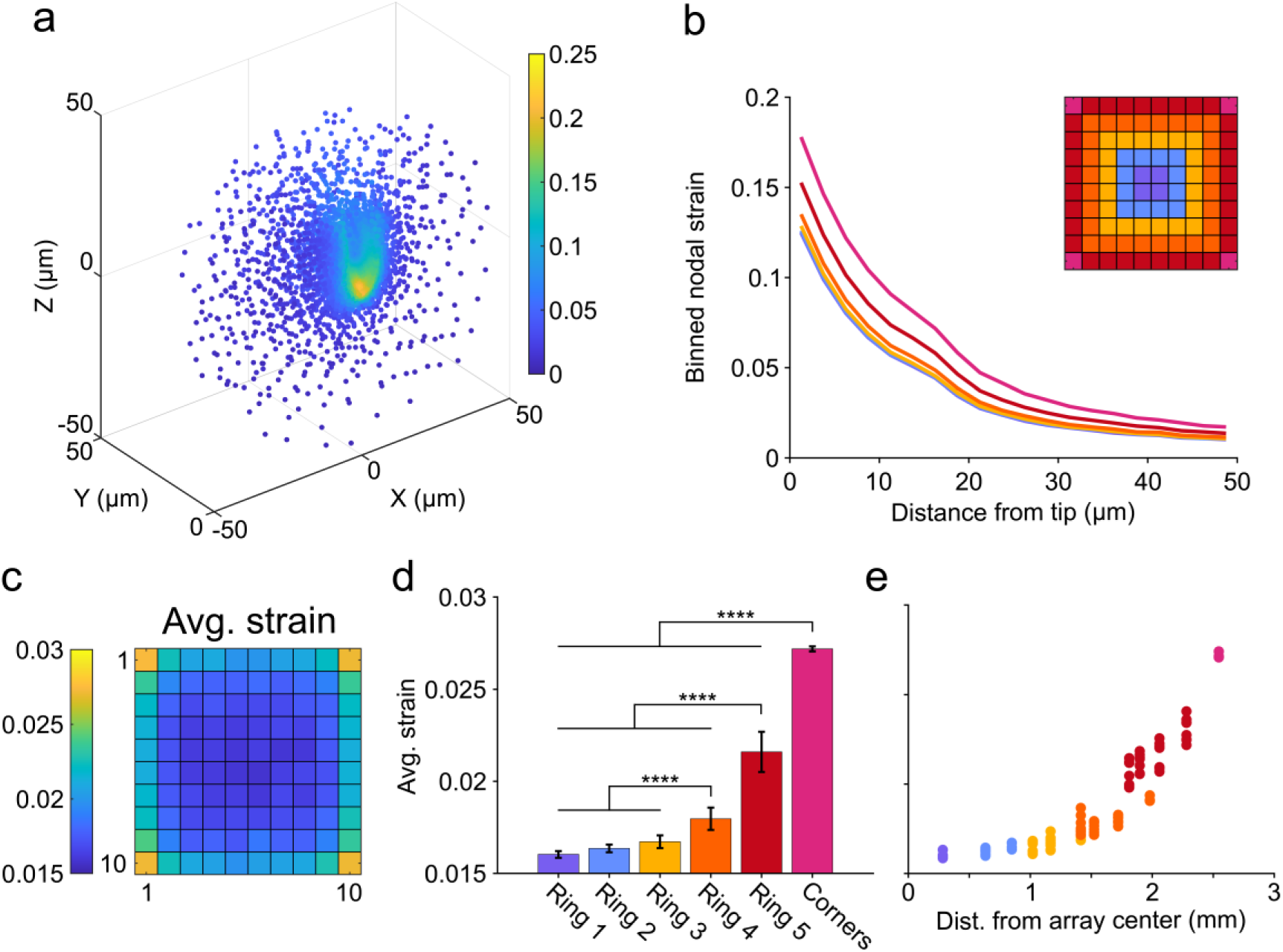
Modeled tissue strains arising from micromotions are greater on edge electrodes compared to interior electrodes. (a) Von Mises strains for brain tissue nodes within 50µm of the electrode tip. Example shown is a corner electrode. (b) Average spatial profile of strain surrounding electrodes in 10x10 array. Inset is color legend defining concentric electrode rings. The corner electrodes are designated as a distinct group. (c) Heatmap of average von Mises strain occurring within 50µm of each electrode. (d) Average strains at corner, ring 5, and ring 4 electrodes were significantly different from all other groups (p<0.0001, one-way ANOVA with Tukey’s post-hoc test). (e) Average strains plotted as a function of distance from the center of the array show corner and ring 5 electrodes are distinctly grouped (right). Error bars are standard deviations. **** p<0.0001.

### 3.2 Modeled tissue strains are negatively correlated with impedance across the lifetime of the implant

Given that electrode placement and array design influence tissue strain, we next examined whether our modeled strain magnitudes were correlated with 1kHz impedance measurements from previously implanted array studies [9, 19, 27, 95, 99]. Recordings were obtained from Utah arrays implanted in human study participants (N = 2 participants, n = 8 arrays) at approximately 1-mo, 1-yr, and 2-yrs post-implantation. The 10x10 arrays (n=4) were implanted in the motor cortex while the 10x6 stimulating arrays (n=4) were implanted in Area 1 of the somatosensory cortex. Similarly, recordings from 8x8 arrays implanted in the V4 region of macaques (N=2 animals, n = 4 arrays) were collected at approximately the same time points.

Impedance values were normalized within individual arrays by subtracting the minimum impedance from all electrodes and then dividing by the maximum. For 10x10 human motor arrays and 8x8 NHP V4 arrays, electrodes near the edge of the array had reduced impedance compared to those in the interior (Kruskal-Wallis test with post-hoc Dunn’s test). Modeled strains were then correlated with the normalized impedances using Spearman’s rank-order correlation. For the 10x10 arrays, a significant negative correlation (p<0.0001) was found for each time period (Fig. 3a-c), with the highest magnitude Spearman’s ρ of -0.39 occurring at 1-yr post-implantation (Fig. 3b); higher strain was associated with lower impedance. No significant correlation was observed for the 10x6 arrays at any time point (Fig. 3d-f). Finally, for the 8x8 arrays, significant negative correlations (p<0.0001) were found at 1-mo and 1-yr after implantation, both having comparable Spearman’s ρ, -0.38 and -0.41, respectively (Fig. 3g-i). These results suggest that impedance can vary with strain, but the relationship is dependent on array geometry and implantation site.

**Figure 3.**
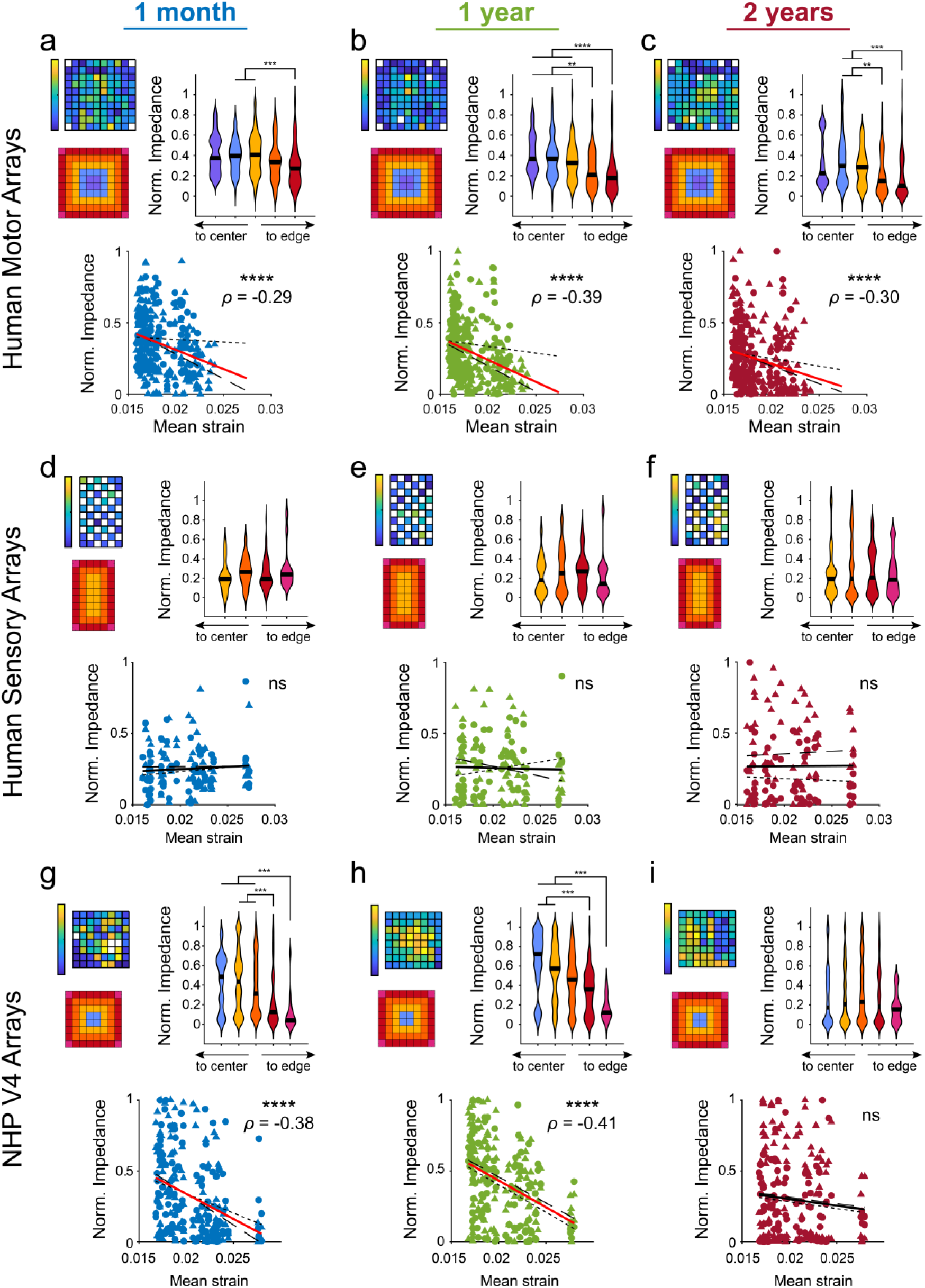
Impedance of implanted Utah arrays are negatively correlated with predicted micromotion strains. (a-c) Normalized 1kHz impedance measured approximately 1-mo, 1-yr, and 2-yrs post-implantation in motor cortex of human study participants (N=2 participants, n=4 arrays). Heatmap shows an example array. Violin plots show edge electrodes had reduced impedances compared to more interior shanks at each time point (Kruskal-Wallis test with post-hoc Dunn’s test). Normalized impedances were correlated with modeled von Mises strains using Spearman’s rank-order correlation. Spearman’s correlation coefficient (ρ) values reported for significant correlations (p<0.05). (d-f) Normalized 1kHz impedance measured approximately 1-mo, 1-yr, and 2-yrs post-implantation in somatosensory cortex of human study participants (N=2 participants, n=4 arrays). Heatmap shows an example array. Violin plots showed impedance did not vary between edge and interior electrodes. (Kruskal-Wallis test with post-hoc Dunn’s test). Normalized impedances were correlated with modeled von Mises strains using Spearman’s rank-order correlation. No significant correlations observed. (g-i) Normalized 1kHz impedance measured approximately 1-mo, 1-yr, and 2-yrs post-implantation in area V4 of macaque monkeys (N=2 animals, n=4 arrays). Heatmap shows an example array. Violin plots show edge electrodes had reduced impedances compared to more interior shanks at 1-mo and 1-yr post-implantation (Kruskal-Wallis test with post-hoc Dunn’s test). Normalized impedances were correlated with modeled von Mises strains using Spearman’s rank-order correlation. Spearman’s correlation coefficient (ρ) values reported for significant correlations (p<0.05). ** p<0.01, *** p<0.001, **** p<0.0001. Black line in violin plots show median value. P2 and Monkey L data is plotted with circles and small-dashed trend line. P3 and Monkey A data is plotted with triangles and large-dashed trend line. The thick trend line fits the combined data.

An alternate hypothesis as to why edge electrodes have reduced impedance compared to interior electrodes could be failure of insulation on the back of the array occurring more frequently on the edge of the array compared to the interior. This would lead to lowered electrical resistance between neighboring electrodes, which could contribute to reduced impedances on individual electrodes. Lower resistance between neighboring electrodes should cause higher correlations in their electrophysiological recordings. To test this alternate hypothesis for the 10x10 human motor arrays, correlations in the bandpass-filtered electrophysiological recordings between neighboring electrodes were computed and then correlated to the collected impedance data (Fig. 4a-c). We found a significant correlation only at 2-yrs post-implantation (Fig. 4c). Therefore, this insulation failure does not explain the reduced edge electrode impedance relative to interior electrodes observed at 1-mo and 1-yr post-implantation. Interestingly, for one 10x10 human motor array, very high electrode neighbor correlations were observed at one edge of the array, and highly correlated to low impedance values (Supplementary Fig. 8, Fig. 4b-c). This suggests that insulation failure at the edge of the array does impact the electrode impedance, but other factors, such as strain, may have larger effects, especially in the first year following implantation.

**Figure 4.**
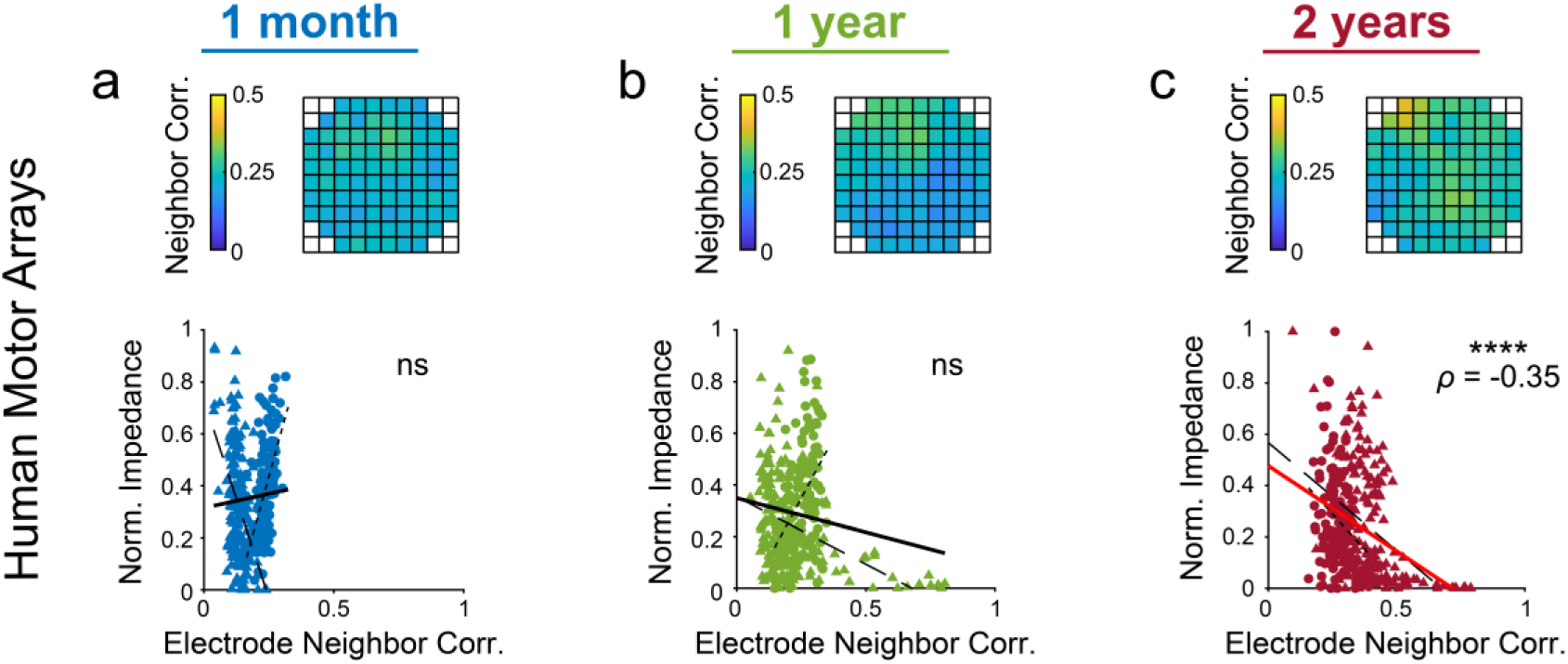
Similarity in bandpass-filtered electrophysiological recordings between neighboring electrodes correlates with impedance at 2-yrs post-implantation in human motor arrays. Using arrays implanted in human motor cortex (N=2 participants, n=4 arrays) at (a) 1-mo, (b) 1-yr, and (c) 2-yrs post-implantation, bandpass-filtered electrophysiological recordings from each electrode were correlated to the bandpass-filtered electrophysiological recordings from all surrounding electrodes. An average correlation was computed for each electrode, estimating its similarity to surrounding electrodes. Heatmap shows an example array. Normalized 1kHz impedance was then correlated with the average neighborhood correlation for that electrode using Spearman’s rank-order correlation. Spearman’s correlation coefficient (ρ) values reported for significant correlations (p<0.05). **** p<0.0001. P2 data is plotted with circles and small-dashed trend line. P3 data is plotted with triangles and large-dashed trend line. The thick trend line fits the combined data.

### 3.3 Modeled tissue strains are negatively correlated with spontaneous PTPV and SNR in human motor arrays years after implantation

Next, we investigated whether our modeled strains were correlated with the quality of recorded spontaneous neural activity from human motor and somatosensory arrays. In the motor cortex, edge electrodes had lower PTPV and SNR compared to interior electrodes at 1-yr and 2-yrs post-implantation (Kruskal-Wallis test with post-hoc Dunn’s test). We also observed a significant negative correlation between strains and both spontaneous PTPV and SNR at 1-yr and 2-yrs post-implantation (p<0.001, Fig. 5b-c, e-f). While insulation failure could explain this trend at 2-yrs post-implantation, the relationship in the first year could be the result of elevated strains reducing the ability of the electrodes from recording nearby neurons. In contrast, no significant correlation between strain and PTPV or SNR was found in the somatosensory arrays. (p>0.05, Supplementary Fig. 9a-f). The limited number of active channels recorded in these arrays may account for the weaker relationships observed. This highlights the complex interplay between mechanical strain, electrode performance, and the longevity of neural recordings.

**Figure 5.**
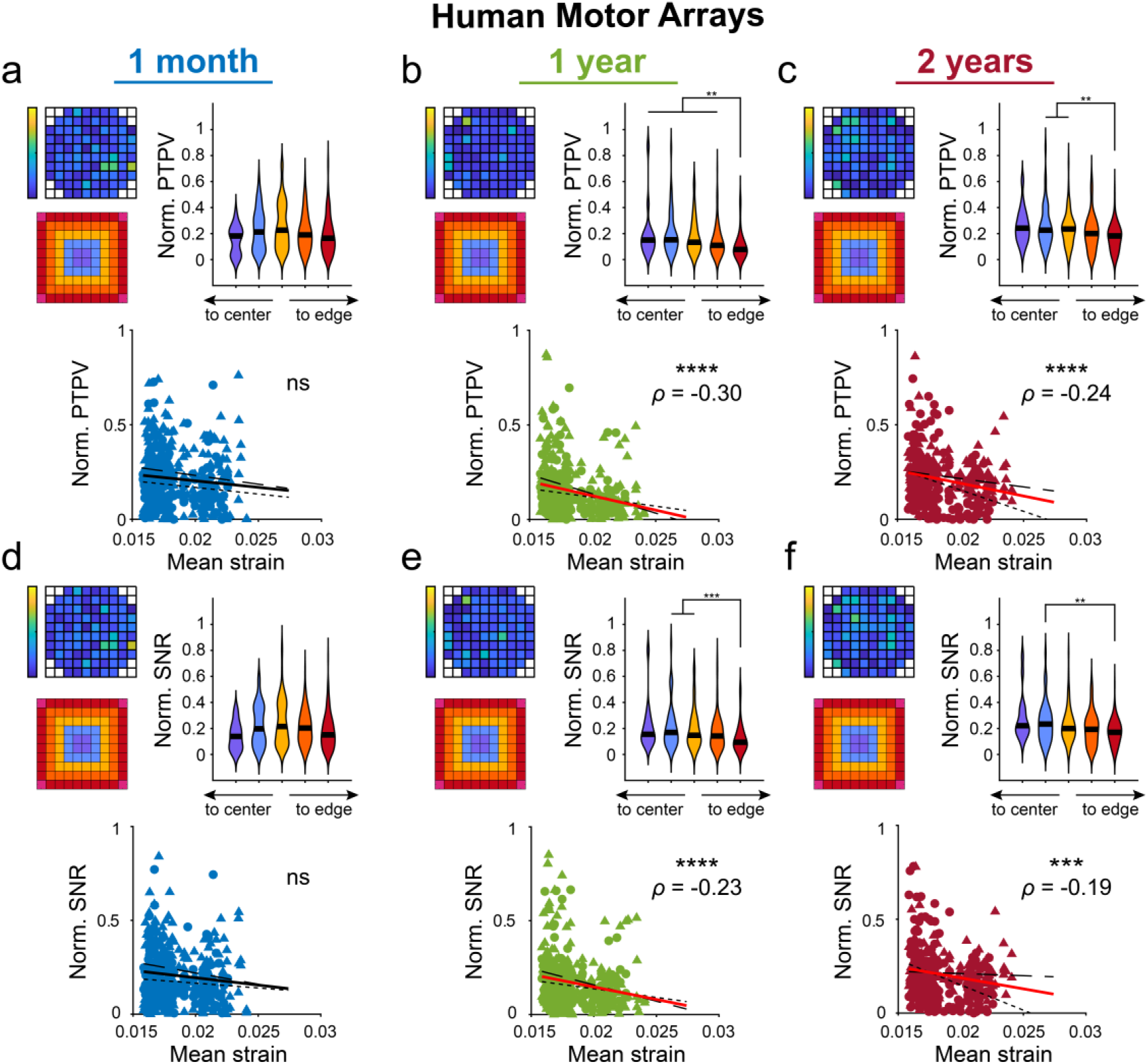
Spontaneous peak-to-peak voltage and SNR from implanted human motor arrays are negatively correlated with predicted micromotion strains. (a-c) Normalized PTPV measured at 1-mo, 1-yr, and 2-yrs post-implantation in motor cortex of human study participants (N=2 participants, n=4 arrays). Heatmap shows an example array. Violin plots show edge electrodes had reduced PTPVs compared to more interior shanks at 1-yr and 2-yrs post-implantation (Kruskal-Wallis test with post-hoc Dunn’s test). Normalized PTPVs were correlated with modeled von Mises strains using Spearman’s rank-order correlation. Spearman’s correlation coefficient (ρ) values reported for significant correlations (p<0.05). (d-f) Same as above, but using normalized SNR measured at 1-mo, 1-yr, and 2-yrs post-implantation in motor cortex of human study participants. Violin plots show edge electrodes had reduced SNRs compared to more interior shanks at 1-yr and 2-yrs post-implantation (Kruskal-Wallis test with post-hoc Dunn’s test). Normalized PTPVs were correlated with modeled von Mises strains using Spearman’s rank-order correlation. Spearman’s correlation coefficient (ρ) values reported for significant correlations (p<0.05). ** p<0.01, *** p<0.001, **** p<0.0001. Black line in violin plots show median value. P2 data is plotted with circles and small-dashed trend line. P3 data is plotted with triangles and large-dashed trend line. The thick trend line fits the combined data.

### 3.4 Modeled tissue strains are positively correlated with envelope MUA-derived evoked SNR in NHP V4 implants

Given that mechanical strain influences electrochemical impedance and spontaneous neural recordings, we next investigated whether predicted tissue strains were correlated with our ability to record visually evoked neuronal activity. Analysis of the PTPV from threshold-crossing snippets showed no correlation with tissue strains in the NHP recordings (Fig. 6a-b). The PTPV data at 2-yrs post-implantation was excluded due to an excessive number of electrodes failing to register any threshold-crossing events. Consequently, we analyzed the envelope MUA and computed the evoked SNR during a stimulus detection task. This analysis revealed a significant positive correlation (p<0.0001 at 1-mo, p<0.01 at 1-yr) between the evoked MUA SNR and tissue strains at 1-mo and 1-yr post-implantation (Fig. 6c-e). Unlike the PTPV results, the positive correlation between MUA SNR and tissue strains suggests that higher mechanical strains are associated with improved evoked SNR, indicating a complex relationship between strain and the quality of visually evoked neural recordings.

**Figure 6.**
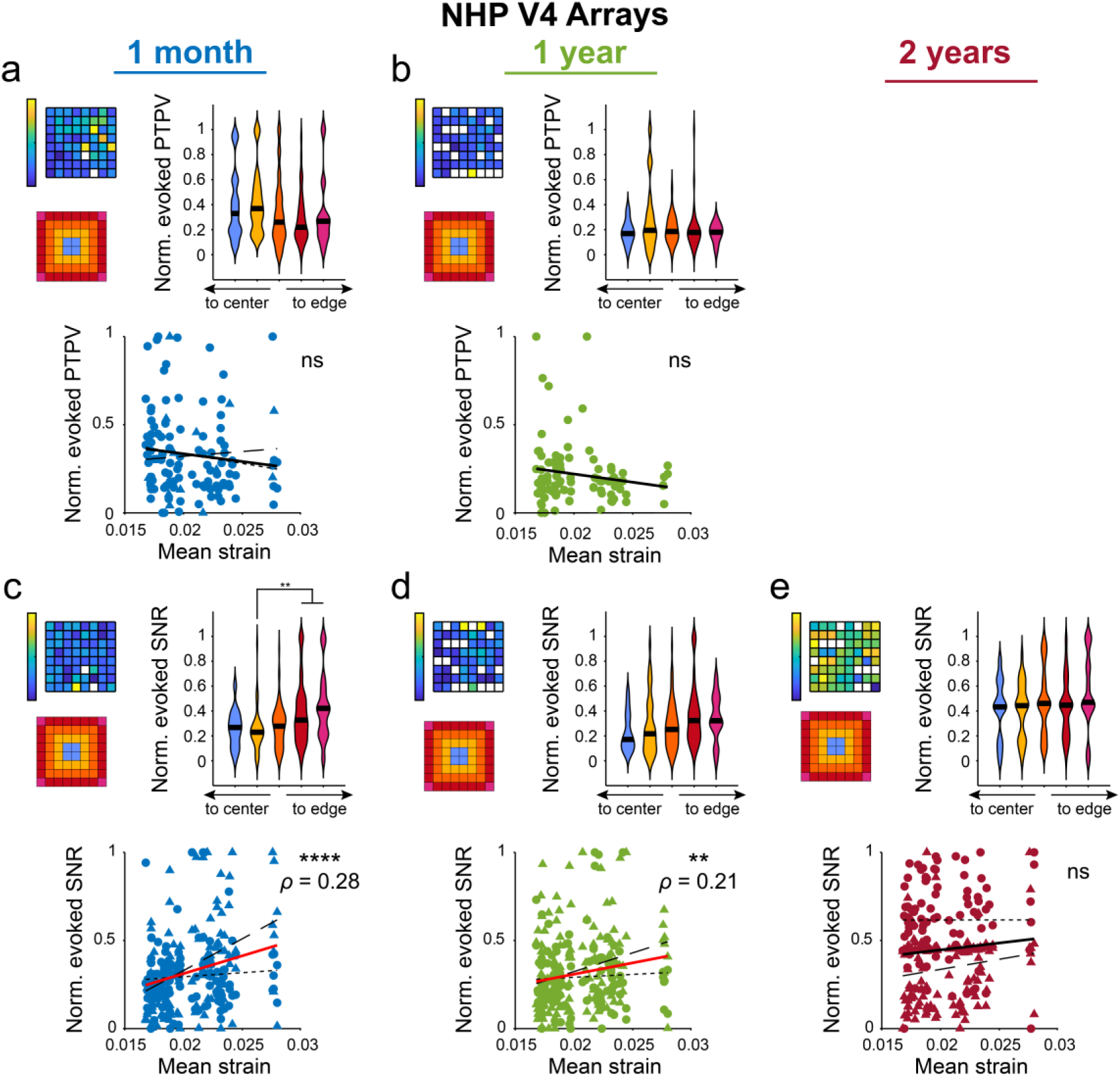
Visually evoked SNR from eMUA recorded from NHP Utah arrays are positively correlated with predicted micromotion strains. (a-b) Normalized PTPV measured at 1-mo and 1-yr post-implantation in area V4 of macaque monkeys (N=2 animals, n=4 arrays). Heatmap shows an example array. Violin plots show PTPV did not vary between edge and interior shanks (Kruskal-Wallis test with post-hoc Dunn’s test). Normalized PTPVs were correlated with modeled von Mises strains using Spearman’s rank-order correlation. No significant correlations observed. (c-e) Normalized SNR from eMUA measured at 1-mo, 1-yr, and 2-yrs post-implantation in NHP V4. Heatmap shows an example array. Violin plots show normalized eMUA SNR was greater on edge electrodes compared to interior shanks at 1-mo post-implantation (Kruskal-Wallis test with post-hoc Dunn’s test). Normalized SNRs were correlated with modeled von Mises strains using Spearman’s rank-order correlation, with significant correlations occurring at 1-mo and 1-yr post-implantation. ** p<0.01, *** p<0.001, **** p<0.0001. Black line in violin plots show median value. Monkey L data is plotted with circles and small-dashed trend line. Monkey A data is plotted with triangles and large-dashed trend line. The thick trend line fits the combined data.

## 4. Discussion

This study aimed to model the strain profiles in tissue surrounding chronically implanted electrodes caused by micromotions and to determine whether these strains correlate with functional metrics from implanted arrays. Utah arrays have been a standard recording modality for human and non-human primate BCIs for decades. However, the impact of array geometry on neuronal recording remains poorly understood. Here, we establish a spatial dependence for tissue strain and array recording capabilities, specifically noting differences between edge and interior shanks. Insights into this relationship are essential for developing improved multi-electrode arrays or placement strategies for serially implanted multi-shank electrodes, with the goal of enhancing longevity and advancing the clinical translation of motor and sensory BCIs.

### 4.1 Signal quality in human motor arrays is negatively correlated with predicted tissue strains

We observed a significant (p<0.001) negative correlation between our modeled tissue strain and the PTPV and SNR from spontaneous recordings in human motor cortex (Fig. 5b-c,e-f). The presence of correlation after 1-yr and lack of correlation at 1-mo may be attributed to increased glial scarring surrounding edge electrodes, a consequence of elevated micromotion-derived strains. Such scarring fosters a neurodegenerative environment near the electrode, diminishing the efficacy of recording and stimulating devices [101–103]. Both microglia and astrocytes can respond to mechanical strains and stiffness gradients in the tissue [43, 53–55, 58, 59, 71–75, 104, 105]. Microglia have been shown to migrate towards stiffer substrates and are more prone to display activated macrophage-like phenotypes in these stiffer environments [55, 104, 106]. For example, in Alzheimer’s Disease models, microglia migration to amyloid beta plaques is mediated by mechanosensitive ion channels [58, 59]. Mechanical stimuli can similarly contribute to astrocyte reactivity via the opening of mechanosensitive ion channels [55, 72, 105]. The heightened strain experienced by corner and edge electrodes (Fig. 2c) likely contributes to increased glial reactivity. Histological studies of implanted Utah arrays support these findings. Notably, Patel et al. reported more frequent damage to edge electrodes compared to interior shanks, aligning with our model’s prediction of greater strain at edge electrodes of an array [107–109].

At early time points, strains exerted on blood vessels near the edge of the Utah array can also contribute to vascular damage and subsequent ischemia in the central regions of the array. High mechanical strain, especially near edge electrodes, can cause significant stress on adjacent blood vessels, leading to endothelial cell damage, vessel rupture, or increased permeability [79, 110]. Such damage can compromise the BBB and result in localized hemorrhage or edema. This vascular injury impairs oxygen and nutrient delivery to the brain tissue, potentially leading to ischemic conditions in downstream regions. The observation of increased strain near edge electrodes (Fig. 2c) could correlate with heightened vascular stress and damage in those areas. Consequently, the compromised blood supply near the edge can affect the central regions of the array, resulting in ischemia. This ischemic environment can severely impact neuronal health and functionality, thereby affecting the quality of neural recordings. For instance, the lack of correlations observed at 1-mo post-implantation (Fig. 5a,d) might be attributed to ischemic damage and silencing neuronal activity in these central regions [52]. However, over the long term, vessels may repair and revascularize the tissue, evidenced by increased PTPV and SNR for central electrodes compared to edge electrodes at 1-yr and 2-yr post-implantation. These findings underscore the impact of mechanical strain on neural recording quality, particularly highlighting how increased strain around edge electrodes correlates with diminished recording performance.

### 4.2 SNR of evoked eMUA in non-human primates is positively correlated with predicted strains

In our analysis of visually evoked recordings from NHP V4, we observed an unexpected positive correlation between predicted strain and the evoked SNR of envelope MUA (Fig. 6c,d). This finding suggests that higher predicted strains are associated with better quality evoked neural signals. The envelope MUA, which aggregates activity from multiple neurons, captures signals from a broader neuronal population than single-unit recordings, potentially from regions beyond the immediate vicinity of the electrode [111–113]. Our model indicates minimal strain variation beyond 40 µm from the electrode tip (Fig. 2b), suggesting that micromotion strains predominantly affect neurons very close to the electrode. Consequently, the observed positive correlation between strain and MUA might reflect that the MUA detects signals from relatively unaffected, healthier tissue further from the electrode. This implies that other factors related to the array geometry or the nature of the evoked response might be influencing the positive correlation.

For instance, the array’s design could enhance signal detection from neurons located in less strained regions or affect the spatial distribution of strain in a way that benefits broader neuronal populations. Additionally, the positive correlation might indicate that the evoked response is more robust in areas with higher strain due to improved coupling between the electrode and surrounding neural tissue, or because the strain could be influencing the electrode-tissue interface in a way that improves signal transmission. Alternatively, the positive correlation observed could be the result of variable tissue properties between human motor cortex and macaque area V4. However, taken together, the positive correlation between predicted strain and MUA in NHP V4 highlights a complex interaction between mechanical strain and the quality of evoked neural recordings, suggesting that higher strain may enhance signal capture from a broader neuronal population, possibly due to improved electrode-tissue coupling or other geometric factors related to the array design.

Also note that a previous study assessing the chronic performance of these NHP arrays showed that by 3 years after implantation, on several arrays, greater tissue encapsulation was observed on electrodes located at the center of the arrays (e.g. Fig. 6b of reference 28 (Chen, 2023)), which may contribute to lower SNR in centrally located electrodes.

### 4.3 Impedance is negatively correlated with predicted tissue strain across the lifetime of the array

Analyzing impedance measurements provides insights into the electrode-tissue interface, revealing how mechanical strains influence electrode performance. We observed a significant negative correlation between impedance and predicted micromotion-derived strains in both human motor and NHP V4 arrays (Fig. 3a-c,g-i). This correlation indicates that as predicted strain increases, impedance decreases, highlighting the impact of mechanical strain on electrode functionality. Interestingly, we did not observe any significant trend when assessing the human S1 arrays (Fig. 3d-f). One explanation is that the rectangular 10x6 geometry limits the number of interior electrodes compared to the 10x10 and 8x8 arrays, which could impact the amount of strain shielding that occurs. Additionally, the reduced number of connected electrodes also limited the amount of data that could be analyzed compared to the human motor and NHP V4 arrays.

Over time, we noted a consistent decrease in impedance, aligning with findings from previous studies [27, 59, 86, 114–116]. This trend may be attributed to increasing fluid accumulation in the peri-electrode space. Elevated fluid levels could result from the mechanical stresses and strains experienced by the electrodes, particularly those located at the edges. This fluid buildup might be due to the gliotic response to the fluid shear stress near the electrode, which is mediated by mechanosensitive ion channels [43]. Mechanosensitive ion channels, such as Piezo1, play a role in detecting mechanical forces and responding to changes in the local environment, including fluid shear stress [43, 54]. The increased micromotion strain around edge electrodes might exacerbate fluid accumulation and enhance the gliotic response, leading to a reduction in impedance. This reduction reflects a deterioration in the electrode-tissue interface, where increased fluid accumulation can disrupt the effective electrical coupling between the electrode and surrounding neural tissue. Additionally, the fluid filled space could alter the transport and diffusion of extracellular species necessary for metabolism [117–121] which could also contribute to varied neuronal health in surrounding tissue and recording capabilities across electrodes.

Increased mechanical strain could also contribute to the mechanical failure of the insulation material surrounding the electrode. The parylene insulation is designed to prevent electrical leakage and ensure stable impedance readings [34, 86, 122]. However, prolonged exposure to mechanical stress and strain can cause micro-cracks or delamination in the insulation layer [122]. These defects can compromise the insulating properties, leading to increased electrical leakage and a subsequent decrease in impedance. As the insulation material degrades, its ability to maintain a stable interface between the electrode and the tissue diminishes, further exacerbating impedance reduction [34]. Taken together, the negative correlation between impedance and predicted strain underscores the relationship between mechanical strain and electrode performance over time. As strain increases, impedance decreases, adversely impacting electrode performance and electrode-tissue interface.

### 4.4 Predicted spatial differences in tissue strain arise from planar geometry and material properties of Utah arrays

Previous work has modeled tissue strain surrounding linear electrode implants [84–90], often focusing on how soft and flexible electrodes could reduce strain compared to traditional silicon probes. These models indicated that soft electrodes might lower strain and scarring [85]. Research on linear probes has also highlighted the importance of electrode site geometry on recording performance [86]. While such probes are common in rodent studies, human and NHP BCI research predominantly uses planar Utah arrays for cortical access. While finite element modeling (FEM) has explored temperature dissipation in these arrays [123, 124], the impact of neighboring shanks in Utah arrays on tissue strain due to micromotion has not been previously assessed.

Similar to previous studies [84, 85], our model used a linear displacement of the arrays with fixed brain tissue to simulate relative motion (Fig. 1b). In an effort to be more consistent with current studies using *ex vivo* mechanical testing of cortical tissue, we applied a nonlinear constitutive model for the brain tissue [92]. Contour plots showed increased von Mises strain at the edges and corners of the array compared to interior electrodes (Fig. 1d). Further analysis indicated that corner electrodes caused greater strain than edge electrodes, which in turn imparted more strain than interior electrodes (Fig. 2c). Across the three array geometries, we found that the average modelled strains were similar (Supplementary Fig. 7). The strain distribution was influenced by the array’s ring structure, with tissue near the edges experiencing higher strain compared to the center (Fig. 2d). This suggests that the Utah array’s planar geometry results in a shielding effect, where the interior tissue experiences less strain.

Modifying the Utah array’s geometry could enhance device performance. For instance, adding a new ring of nonfunctional shanks might improve strain shielding but could also increase cortical damage and insertion difficulty. Additionally, using softer or more flexible materials, as seen with linear arrays, could reduce mechanical mismatch and glial scarring [61–68, 125, 126]. Developing free-floating wireless arrays might also minimize tangential tethering forces, potentially reducing tissue strains [127]. New coatings could improve integration with the extracellular matrix and alter how forces affect nearby cells [128]. Taken together, future designs of planar arrays should therefore consider how changes in geometry and material properties impact the strain experienced by surrounding tissues.

### 4.5 Limitations

Many factors could influence the observed correlations, and this study does not establish tissue strain as a causal factor for array performance. Instead, it highlights how Utah array performance and the surrounding cellular environment vary with array geometry. Other contributing factors may include the metabolic support of neurons, which is crucial for survival near the electrode-tissue interface. Neurons on edge and corner electrodes might benefit from proximity to less damaged tissue, enhancing their survival compared to those in the array’s center.

The curvature of the brain could also affect how array geometry impacts recording performance. For electrocorticography grids, brain curvature influences the forces applied to cortical tissue [129]. A similar effect may occur with Utah arrays, where shank orientation relative to the brain surface could impact strain distribution. Additionally, the laminar structure of the cortex might lead to differences in neuronal layers recorded by edge versus interior electrodes, potentially contributing to the observed ring-structured performance correlations (Fig. 5b-c, e-f) [130, 131].

Insertion methods also play a role as demonstrated by linear arrays studies [29, 48, 76, 77, 80, 132]. Unlike linear arrays, Utah arrays are typically implanted using a pneumatic inserter, which introduces strain differently [133, 134]. A recent study showed insertion velocity affects strain near the tip but not along the shank [135]. Although this study did not assess array geometry, similar strain distributions might occur during insertion as observed with micromotions (Fig. 2c), potentially impacting performance.

Differences in recording conditions and surgical procedures between the human and NHP experiments limit our ability to draw clear comparisons between the arrays as many confounds may impact the observed correlations. These comparisons are further restricted by our limited sample size. Finally, this modeling study made critical assumptions about the array’s material properties and applied forces. It also did not account for insertion-induced strains, assuming initial tissue strain was zero. Despite these limitations, the modeling provides a useful approximation, emphasizing that tissue conditions vary across array geometries and correlate with array performance.

## 5. Conclusion

Utah arrays are robust tools for recording neurons in intracortical BCI applications. Here, we establish how strain fields and recording performance can vary across multi-shank microelectrode arrays. Our FEM shows that tissue near edge and corner electrodes experience greater micromotion-derived strains compared to interior electrodes. These predicted strains are negatively correlated with impedances measured 1-mo, 1-yr, and 2-yrs after implantation. Additionally, strains are negatively correlated with SNR and PTPV of spontaneous neural activity recorded in human motor cortex while positively correlated with evoked SNR using eMUA from recordings in NHP area V4. Although tissue strain appears to be a significant factor affecting recording quality, other factors such as metabolic support may also play a role. Overall, our findings underscore that the geometry of the Utah array influences both electrode impedance and neural recording capabilities.

## Acknowledgements

The authors would like to thank Christopher Hughes, Pieter Roelfsema, X. Tracy Cui, Andy Schwartz, and Delin Shi for their advice and input in addition to their willingness to share their neural recording data. We also graciously thank the clinical study participants for their commitment and effort. This work was supported by: NIH R01NS105691, NIH R01NS115707, NIH R01NS129632, NIH R01DC011311, NSF CBET CAREER 1943906, the Defense Advanced Research Projects Agency (DARPA) and Space and Naval Warfare Systems Center Pacific (SSC Pacific) under Contracts N66001-10-C-4056 and N66001-16-C-4051, the National Institute Of Neurological Disorders And Stroke of the National Institutes of Health under Award Numbers UH3NS107714 and R01NS121079, NWO (STW Grant Number P15-42 ‘NESTOR’; ALW Grant Number 823-02-010 and Cross-over Grant Number 17619 ‘INTENSE’), the European Union (ERC Grant Numbers 339490 ‘Cortic_al_gorithms’ and 101052963 ‘NUMEROUS,’ H2020 Research and Innovation programme Grant Number 899287 ‘NeuraViper’), Human Brain Project (Grant Number 650003), NIH DP2EY037405, NIH P30 EY08098, and Research to Prevent Blindness RK Mellon Foundation. Any opinions, findings and conclusions or recommendations expressed in this material are those of the authors and do not necessarily reflect the views of DARPA, SSC Pacific, or the National Institutes of Health.

## CRediT Authorship Statement

**Adam M. Forrest:** Conceptualization, Methodology, Project administration, Data curation, Formal analysis, Visualization, Writing - Original Draft, Writing - Review & Editing. **Nicolas G. Kunigk:** Data curation, Formal analysis, Writing - Review & Editing. **Jennifer L. Collinger:** Methodology, Funding acquisition, Writing - Review & Editing. **Robert A. Gaunt:** Methodology, Funding acquisition, Writing - Review & Editing. **Xing Chen:** Methodology, Data curation, Formal analysis, Funding acquisition, Writing - Review & Editing. **Jonathan P. Vande Geest:** Conceptualization, Methodology, Supervision, Funding acquisition, Writing - Review & Editing. **Takashi D.Y. Kozai:** Conceptualization, Methodology, Supervision, Funding acquisition, Writing - Review & Editing

---

**Supplementary Figure 1.**
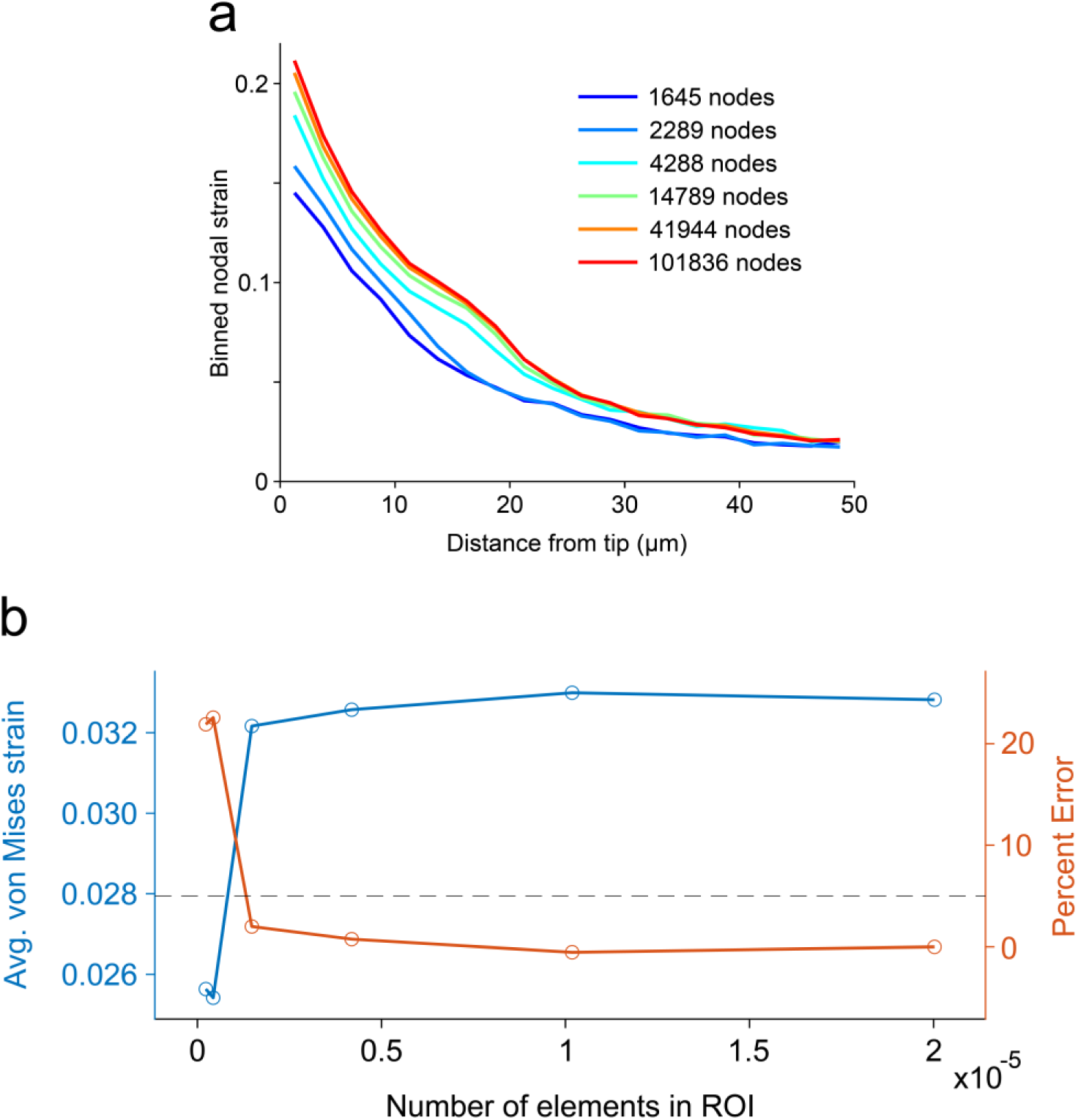
Convergence study to determine finite element modeling mesh size. (a) Convergence study was conducted on a single electrode to reduce computation time. Element size surrounding the electrode tip was refined to achieve higher density meshes. Nodal strains were averaged in 2.5µm bins surrounding electrode tip. (b) Average von Mises strain, computed using the binned strains in (a), converges with increasing element number. The meshing parameters for the third mesh (with 4288 nodes) were selected as an optimal tradeoff of accuracy and computational time.

**Supplementary Figure 2.**
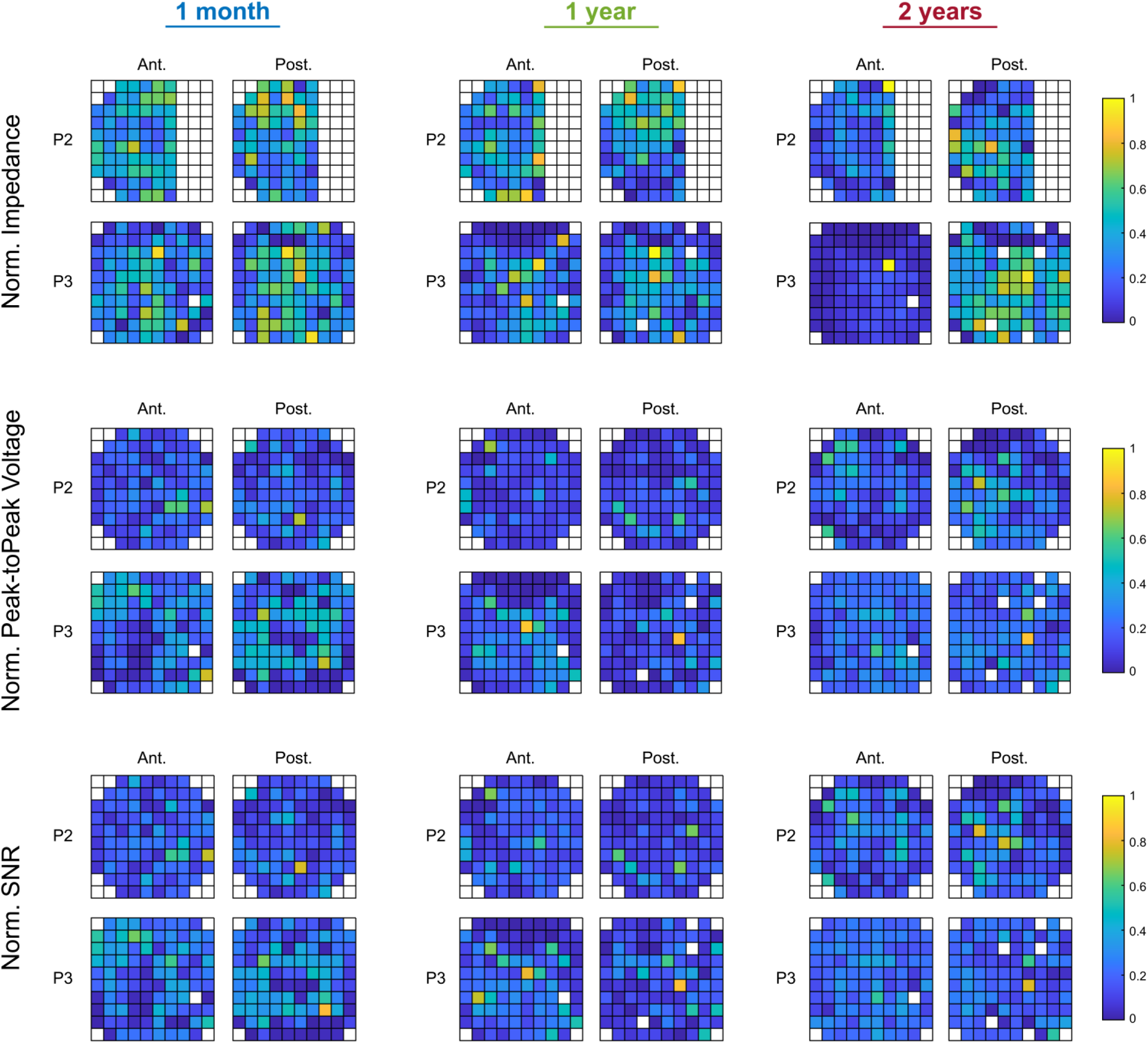
Human motor array performance metrics. For each time period of interest (1-mo, 1-yr, and 2-yrs post-implantation) a total of five successive recording sessions were analyzed (N = 2 participants, n = 4 arrays). For each array, the performance measures were first averaged across the five recording sessions and then normalized within each array. Heatmaps show the normalized performance metrics for each array. Electrodes whited out were either not used for recordings or had 1kHz impedance greater than 1MΩ on every recording day. The large bank of electrodes missing impedance data from P2 were used for recording, but impedance measurements were not collected.

**Supplementary Figure 3.**
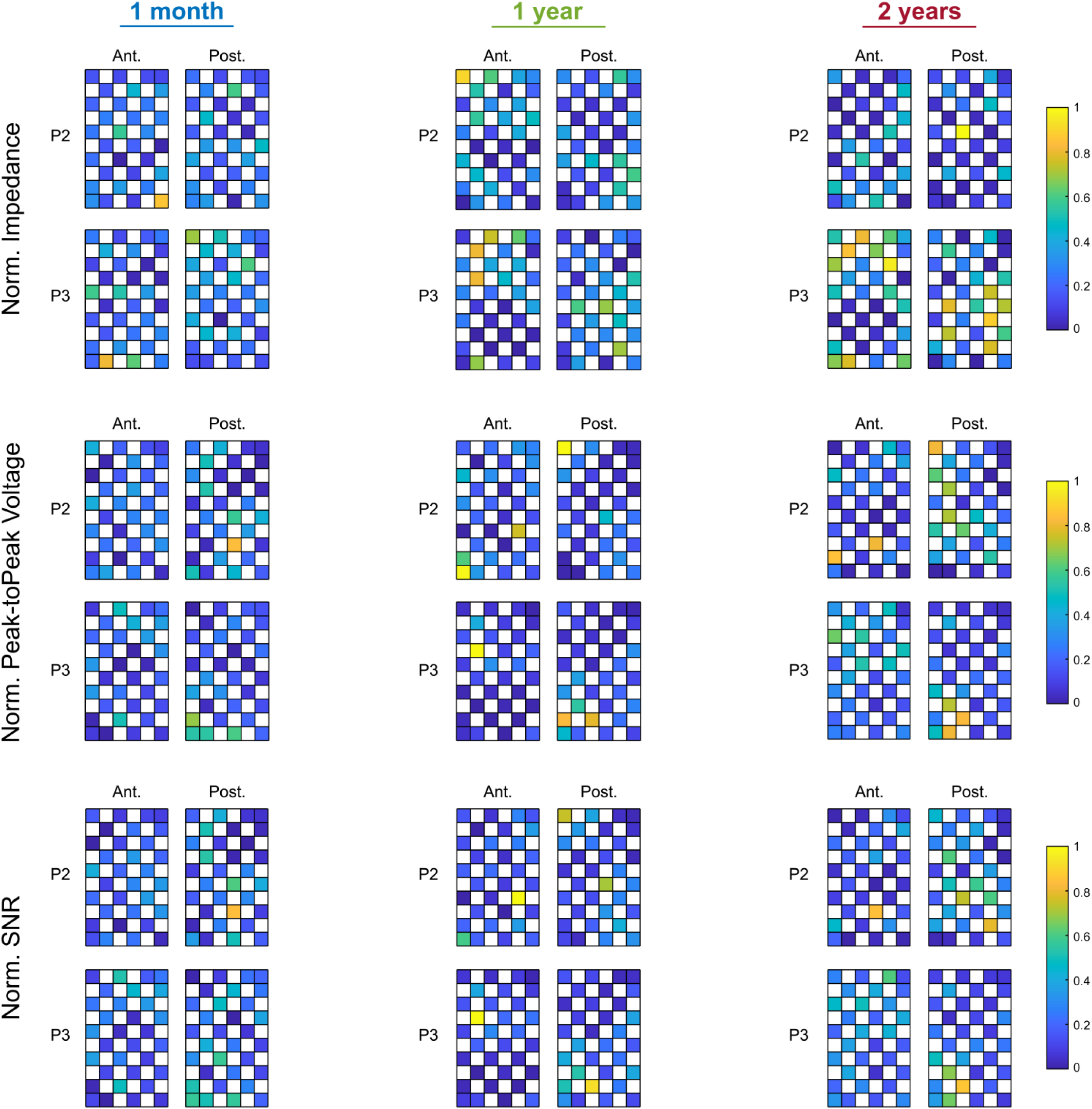
Human somatosensory array performance metrics. For each time period of interest (1-mo, 1-yr, and 2-yrs post-implantation) a total of five successive recording sessions were analyzed (N = 2 participants, n = 4 arrays). For each array, the performance measures were first averaged across the five recording sessions and then normalized within each array. Heatmaps show the normalized performance metrics for each array. Electrodes whited out were not used for recordings.

**Supplementary Figure 4.**
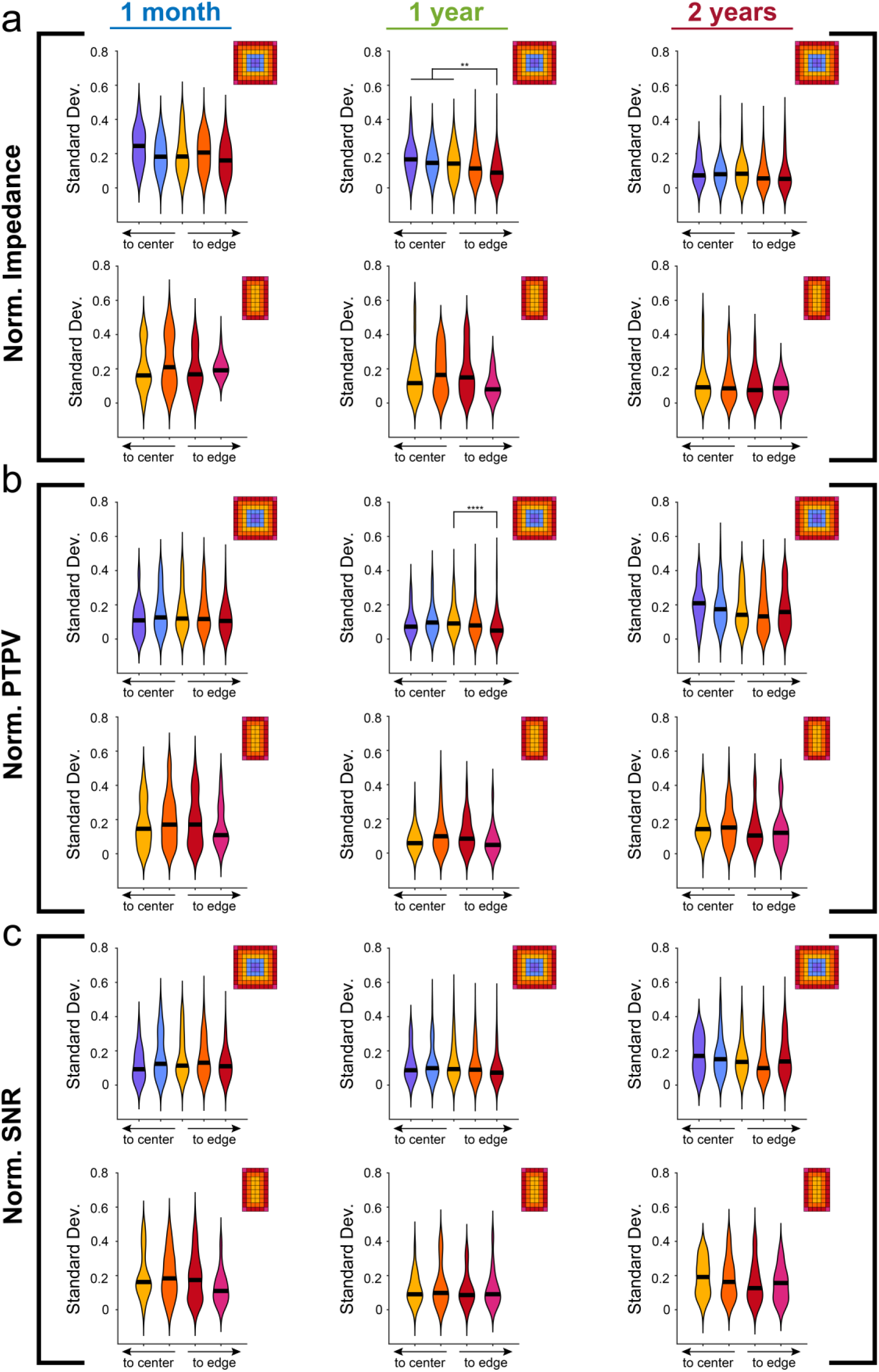
Session-to-session variability shows some dependence on electrode location within the array. Violin plots represent standard deviations of (a) 1kHz impedance, (b) PTPV, and (c) SNR for each electrode across five successive recording sessions. For impedance and SNR metrics, electrodes whited-out are from electrodes whose 1kHz impedance was greater than 1MΩ. No significant differences in session-to-session variability between edge and interior electrodes were observed for the 10x6 somatosensory arrays (Kruskal-Wallis test with post-hoc Dunn’s test). For 10x10 motor arrays, variability in the impedance and PTPV of edge electrodes 1-yr post-implantation was significantly lower compared to interior electrodes (Kruskal-Wallis test with post-hoc Dunn’s test). ** p<0.01, *** p<0.001, **** p<0.0001.

**Supplementary Figure 5.**
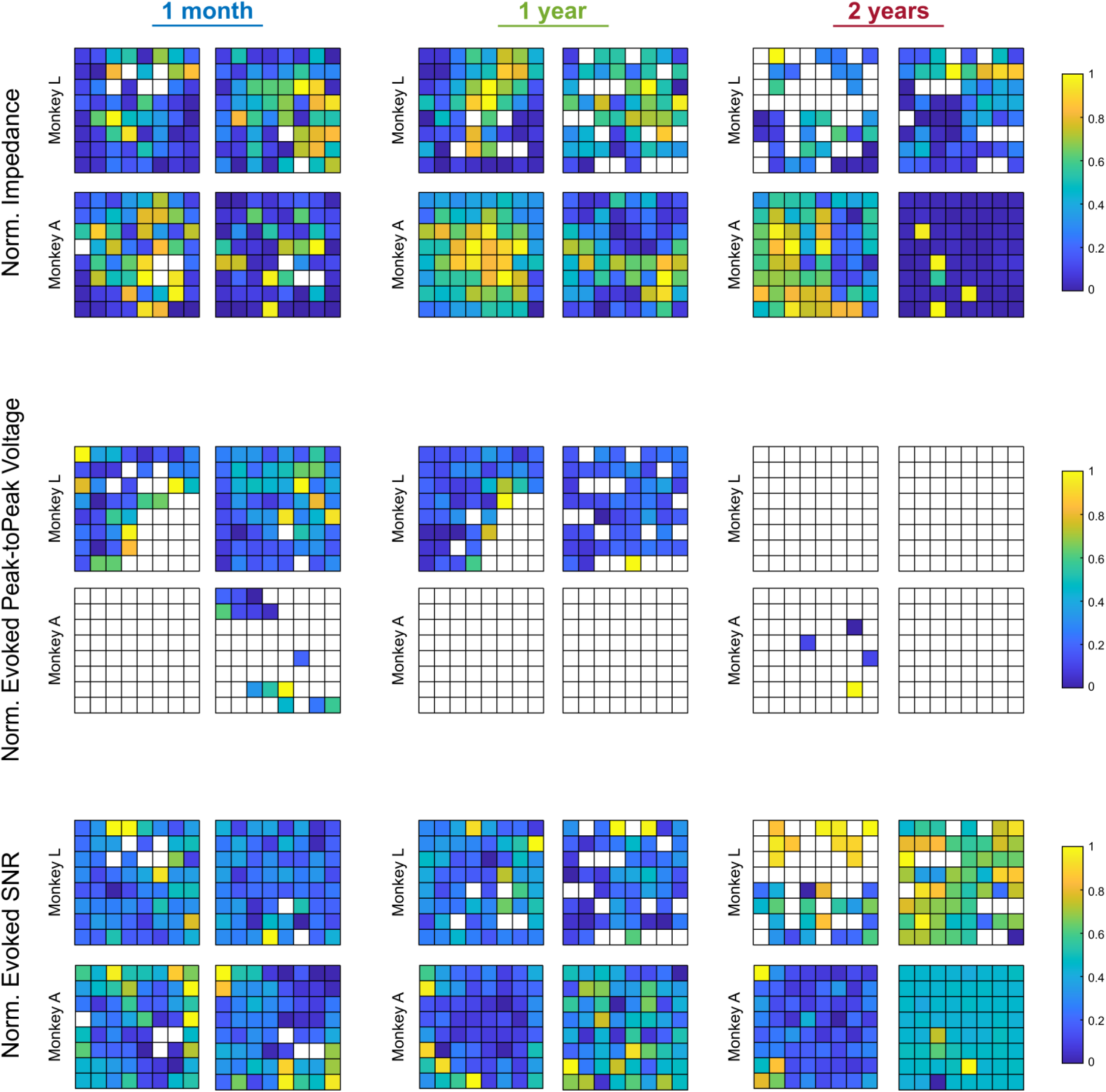
Non-human primate area V4 array performance metrics. Heatmaps show normalized performance measures for each array (N = 2 animals, n = 4 arrays). Limited number of recording sessions prevented analysis for session-to-session variability. Electrodes whited out either had 1kHz impedance greater than 1MΩ or represent electrodes with very low PTPV amplitudes (less than 30µV) and were excluded from the PTPV analysis.

**Supplementary Figure 6.**
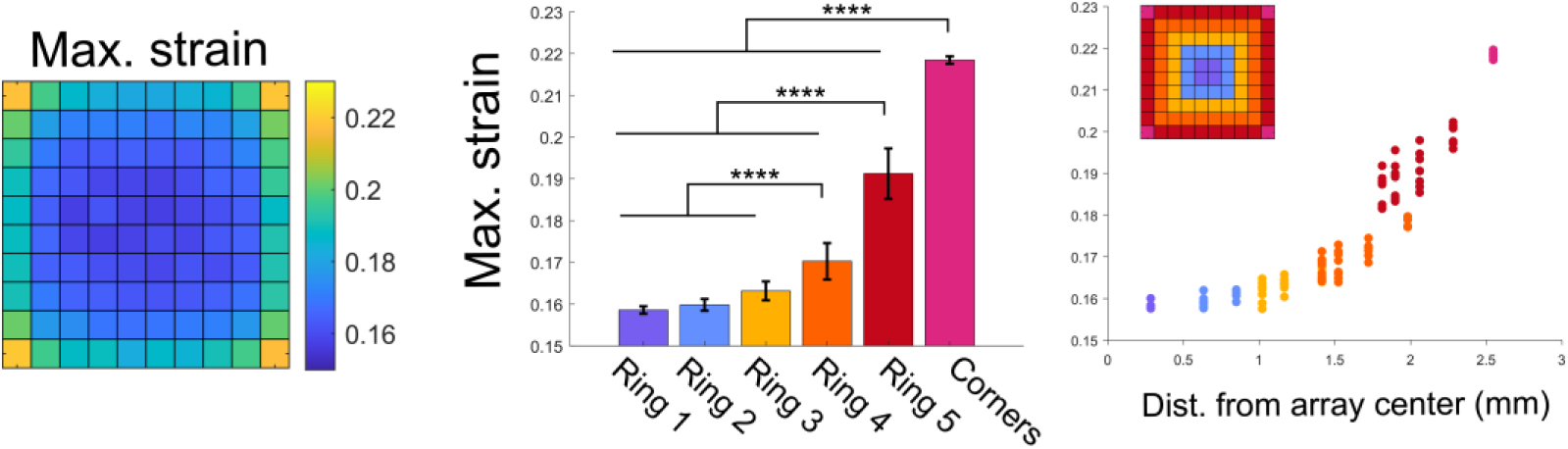
Predicted maximal von Mises strains for 10x10 array geometry show similar trends as average strains. Heatmap of maximum von Mises strain occurring within 50µm of each electrode for 10x10 array (left). Maximum strains at corner, ring 5, and ring 4 electrodes were significantly different from all other groups (p<0.0001, one-way ANOVA with Tukey’s post-hoc test). Maximum strains plotted as a function of distance from the center of the array show corner and ring 5 electrodes are distinctly grouped (right). Error bars are standard deviations. ** p<0.01, *** p<0.001, **** p<0.0001.

**Supplementary Figure 7.**
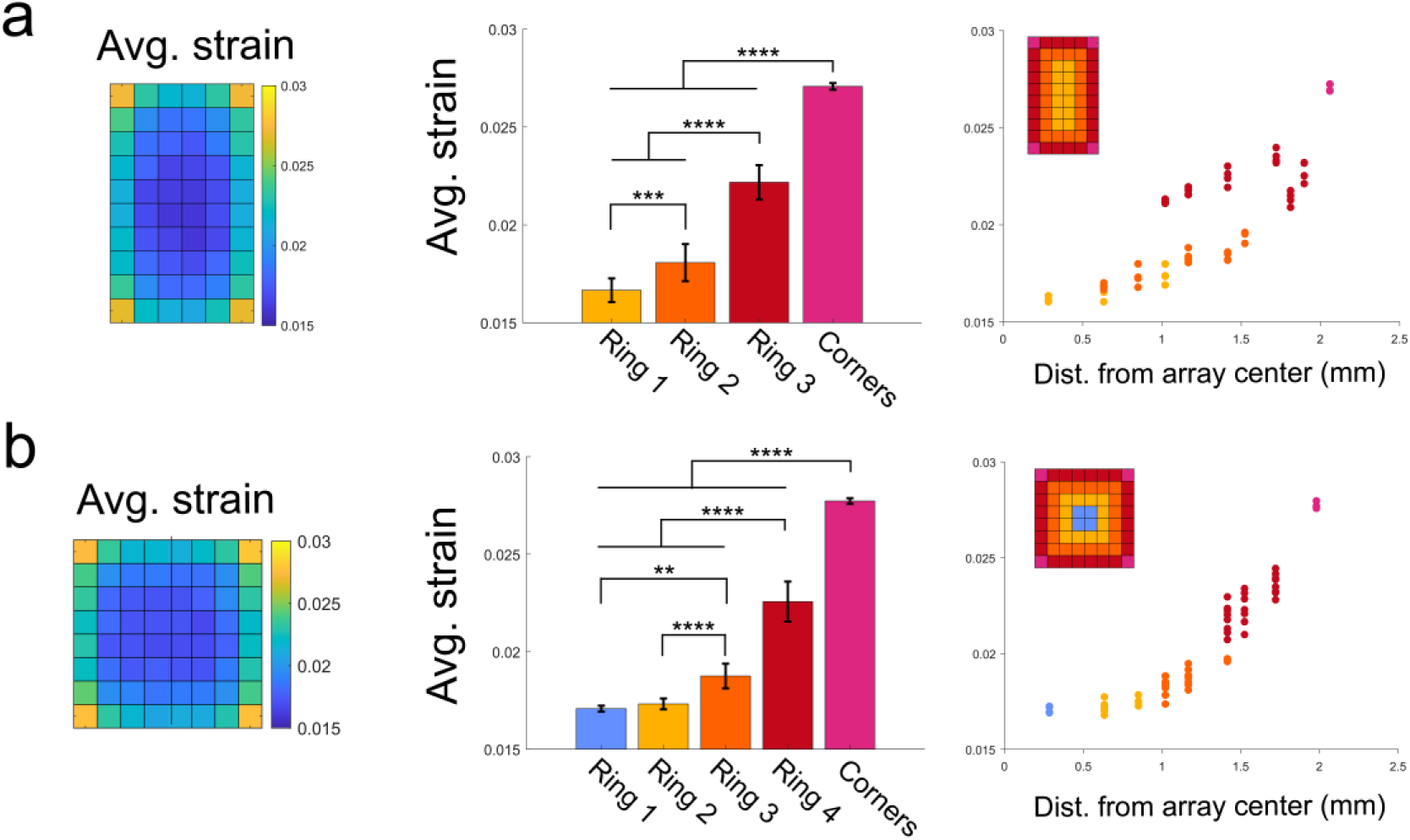
Predicted average von Mises strains for 10x6 and 8x8 array geometries are similar to 10x10 geometry. (a) Heatmap of average von Mises strain occurring within 50µm of each electrode for 10x6 array (left). Average strains for each group were significantly different from all other groups (p<0.001, one-way ANOVA with Tukey’s post-hoc test). Average strains plotted as a function of distance from the center of the array show corner and ring 3 electrodes are distinctly grouped (right). (b) Heatmap of average von Mises strain occurring within 50µm of each electrode for 8x8 array (left). Average strains at corner, ring 4, and ring 3 electrodes were significantly different from all other groups (p<0.01, one-way ANOVA with Tukey’s post-hoc test). Average strains plotted as a function of distance from the center of the array show corner and ring 4 electrodes are distinctly grouped (right). Error bars are standard deviations. ** p<0.01, *** p<0.001, **** p<0.0001.

**Supplementary Figure 8.**
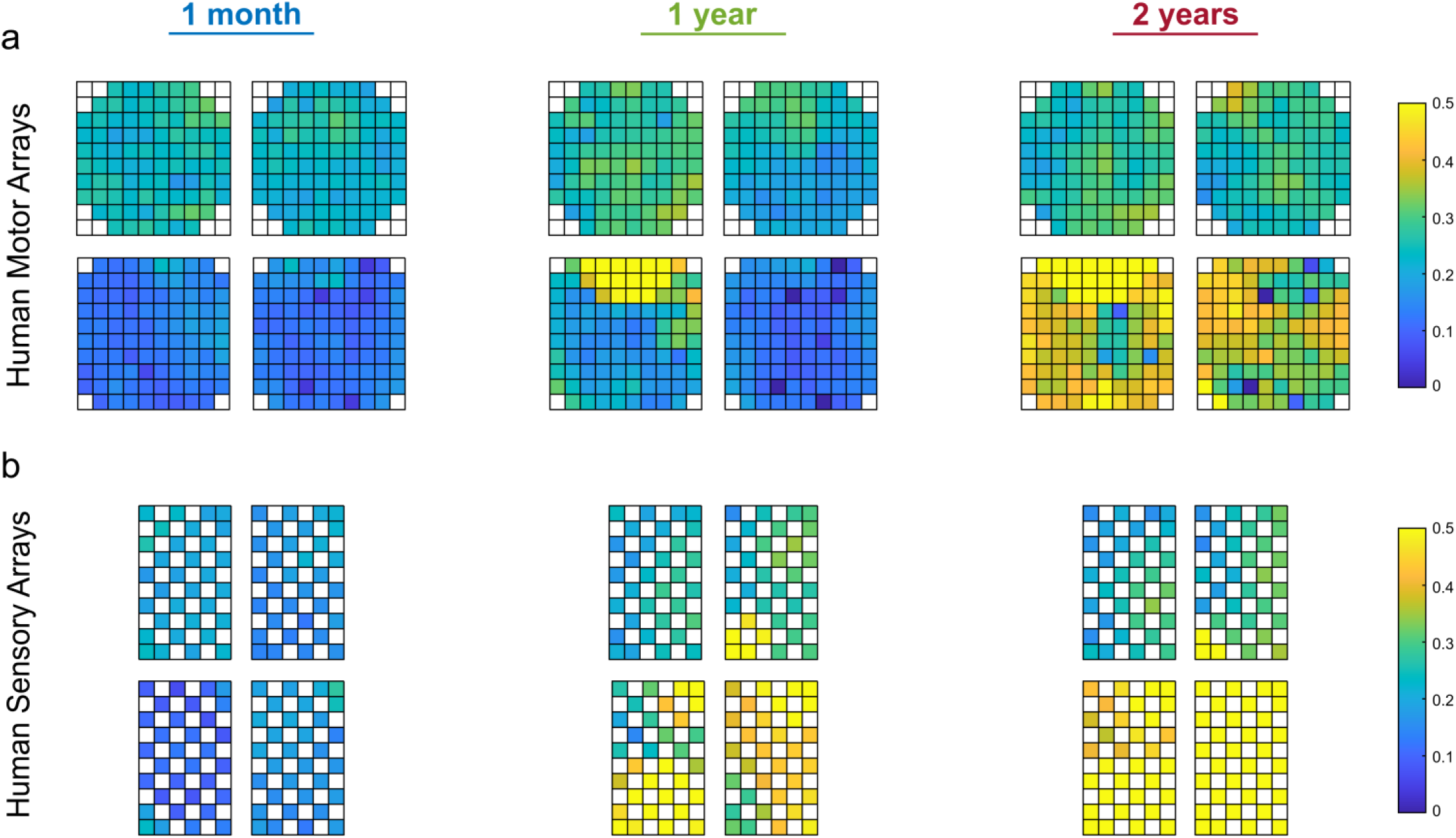
Average correlation with neighboring bandpass-filtered electrophysiological recordings. Heatmaps for (a) 10x10 human motor arrays and (b) 10x6 human somatosensory arrays showing correlation between the bandpass-filtered electrophysiological recordings of neighboring electrodes for all arrays and studied timepoints. Briefly, for each electrode, its bandpass-filtered recording was correlated with 3-8 neighboring electrodes (depending on electrode location). The average of these 3-8 neighboring electrode correlations was then computed. This was repeated for five successive recording sessions for each time period. Heatmaps shown are averages across the five successive recording sessions and whose values are correlated with impedance in Figure 4.

**Supplementary Figure 9.**
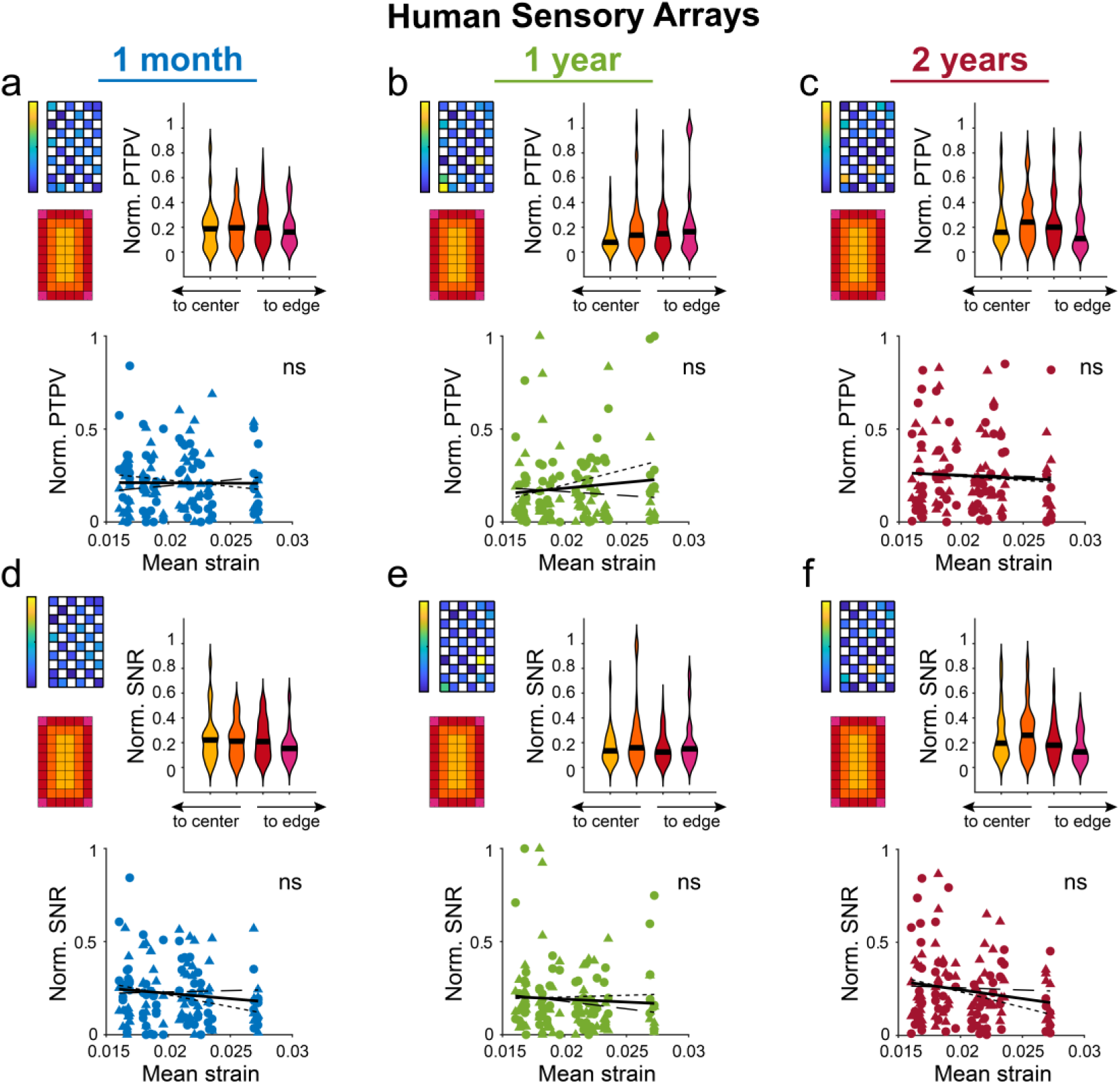
Spontaneous peak-to-peak voltage and SNR from implanted human somatosensory arrays show limited relationship with micromotion strains. (a-c) Normalized PTPV measured at 1-mo, 1-yr, and 2-yrs post-implantation in somatosensory cortex of human study participants (N=2 participants, n=4 arrays). Heatmap shows an example array. Violin plots show no dependence on electrode location within the array (Kruskal-Wallis test with post-hoc Dunn’s test). Normalized PTPVs were correlated with modeled von Mises strains using Spearman’s rank-order correlation. No significant correlation as observed. (d-f) Same as above, but using normalized SNR measured at 1-mo, 1-yr, and 2-yrs post-implantation in somatosensory cortex of human study participants. No strain correlation or edge/interior dependence was observed. Black line in violin plots show median value. P2 data is plotted with circles and small-dashed trend line. P3 data is plotted with triangles and large-dashed trend line. The thick trend line fits the combined data.

## References

[1] N.G. Hatsopoulos, J.P. Donoghue, The science of neural interface systems, Annu Rev Neurosci 32 (2009) 249–66.

[2] M. Bamdad, H. Zarshenas, M.A. Auais, Application of BCI systems in neurorehabilitation: a scoping review, Disabil Rehabil Assist Technol 10(5) (2015) 355–64.

[3] A.B. Schwartz, D.M. Taylor, S.I. Tillery, Extraction algorithms for cortical control of arm prosthetics, Curr Opin Neurobiol 11(6) (2001) 701–7.

[4] L.R. Hochberg, M.D. Serruya, G.M. Friehs, J.A. Mukand, M. Saleh, A.H. Caplan, A. Branner, D. Chen, R.D. Penn, J.P. Donoghue, Neuronal ensemble control of prosthetic devices by a human with tetraplegia, Nature 442(7099) (2006) 164- 71.

[5] L.R. Hochberg, D. Bacher, B. Jarosiewicz, N.Y. Masse, J.D. Simeral, J. Vogel, S. Haddadin, J. Liu, S.S. Cash, P. van der Smagt, J.P. Donoghue, Reach and grasp by people with tetraplegia using a neurally controlled robotic arm, Nature 485(7398) (2012) 372-5.

[6] J.L. Collinger, B. Wodlinger, J.E. Downey, W. Wang, E.C. Tyler-Kabara, D.J. Weber, A.J. McMorland, M. Velliste, M.L. Boninger, A.B. Schwartz, High-performance neuroprosthetic control by an individual with tetraplegia, Lancet 381(9866) (2013) 557-64.

[7] B. Wodlinger, J.E. Downey, E.C. Tyler-Kabara, A.B. Schwartz, M.L. Boninger, J.L. Collinger, Ten-dimensional anthropomorphic arm control in a human brain-machine interface: difficulties, solutions, and limitations, J Neural Eng 12(1) (2015) 016011.

[8] R.A. Parker, T.S. Davis, P.A. House, R.A. Normann, B. Greger, The functional consequences of chronic, physiologically effective intracortical microstimulation, Prog Brain Res 194 (2011) 145–65.

[9] S.N. Flesher, J.L. Collinger, S.T. Foldes, J.M. Weiss, J.E. Downey, E.C. Tyler-Kabara, S.J. Bensmaia, A.B. Schwartz, M.L. Boninger, R.A. Gaunt, Intracortical microstimulation of human somatosensory cortex, Sci Transl Med 8(361) (2016) 361ra141.

[10] M. Armenta Salas, L. Bashford, S. Kellis, M. Jafari, H. Jo, D. Kramer, K. Shanfield, K. Pejsa, B. Lee, C.Y. Liu, R.A. Andersen, Proprioceptive and cutaneous sensations in humans elicited by intracortical microstimulation, Elife 7 (2018).

[11] E. Fernandez, A. Alfaro, C. Soto-Sanchez, P. Gonzalez-Lopez, A.M. Lozano, S. Pena, M.D. Grima, A. Rodil, B. Gomez, X. Chen, P.R. Roelfsema, J.D. Rolston, T.S. Davis, R.A. Normann, Visual percepts evoked with an intracortical 96-channel microelectrode array inserted in human occipital cortex, J Clin Invest 131(23) (2021).

[12] F. Grani, C. Soto-Sanchez, F.D. Farfan, A. Alfaro, M.D. Grima, A. Rodil Doblado, E. Fernandez, Time stability and connectivity analysis with an intracortical 96-channel microelectrode array inserted in human visual cortex, J Neural Eng 19(4) (2022).

[13] L.E. Osborn, B.P. Christie, D.P. McMullen, R.W. Nickl, M.C. Thompson, A.S. Pawar, T.M. Thomas, M. Alejandro Anaya, N.E. Crone, B.A. Wester, S.J. Bensmaia, P.A. Celnik, G.L. Cantarero, F.V. Tenore, M.S. Fifer, Intracortical microstimulation of somatosensory cortex enables object identification through perceived sensations, Annu Int Conf IEEE Eng Med Biol Soc 2021 (2021) 6259–6262.

[14] R.J. Vetter, J.C. Williams, J.F. Hetke, E.A. Nunamaker, D.R. Kipke, Chronic neural recording using silicon-substrate microelectrode arrays implanted in cerebral cortex, IEEE Trans Biomed Eng 51(6) (2004) 896–904.

[15] C.A. Chestek, V. Gilja, P. Nuyujukian, J.D. Foster, J.M. Fan, M.T. Kaufman, M.M. Churchland, Z. Rivera-Alvidrez, J.P. Cunningham, S.I. Ryu, K.V. Shenoy, Long-term stability of neural prosthetic control signals from silicon cortical arrays in rhesus macaque motor cortex, J Neural Eng 8(4) (2011) 045005.

[16] G.W. Fraser, A.B. Schwartz, Recording from the same neurons chronically in motor cortex, J Neurophysiol 107(7) (2012) 1970–8.

[17] J.E. Downey, N. Schwed, S.M. Chase, A.B. Schwartz, J.L. Collinger, Intracortical recording stability in human brain-computer interface users, J Neural Eng 15(4) (2018) 046016.

[18] A.J. Bullard, B.C. Hutchison, J. Lee, C.A. Chestek, P.G. Patil, Estimating Risk for Future Intracranial, Fully Implanted, Modular Neuroprosthetic Systems: A Systematic Review of Hardware Complications in Clinical Deep Brain Stimulation and Experimental Human Intracortical Arrays, Neuromodulation 23(4) (2020) 411–426.

[19] C.L. Hughes, S.N. Flesher, J.M. Weiss, J.E. Downey, M. Boninger, J.L. Collinger, R.A. Gaunt, Neural stimulation and recording performance in human sensorimotor cortex over 1500 days, Journal of Neural Engineering 18(4) (2021) 045012.

[20] Y. Wang, X. Yang, X. Zhang, Y. Wang, W. Pei, Implantable intracortical microelectrodes: reviewing the present with a focus on the future, Microsyst Nanoeng 9 (2023) 7.

[21] K.M. Patrick-Krueger, I. Burkhart, J.L. Contreras-Vidal, The state of clinical trials of implantable brain–computer interfaces, Nature Reviews Bioengineering (2024).

[22] P.J. Rousche, R.A. Normann, Chronic recording capability of the Utah Intracortical Electrode Array in cat sensory cortex, Journal of neuroscience methods 82(1) (1998) 1–15.

[23] J.C. Williams, R.L. Rennaker, D.R. Kipke, Long-term neural recording characteristics of wire microelectrode arrays implanted in cerebral cortex, Brain Res Brain Res Protoc 4(3) (1999) 303–13.

[24] J.C. Barrese, N. Rao, K. Paroo, C. Triebwasser, C. Vargas-Irwin, L. Franquemont, J.P. Donoghue, Failure mode analysis of silicon-based intracortical microelectrode arrays in non-human primates, J Neural Eng 10(6) (2013) 066014.

[25] C. Sponheim, V. Papadourakis, J.L. Collinger, J. Downey, J. Weiss, L. Pentousi, K. Elliott, N.G. Hatsopoulos, Longevity and reliability of chronic unit recordings using the Utah, intracortical multi-electrode arrays, J Neural Eng 18(6) (2021).

[26] L.J. Szymanski, S. Kellis, C.Y. Liu, K.T. Jones, R.A. Andersen, D. Commins, B. Lee, D.B. McCreery, C.A. Miller, Neuropathological effects of chronically implanted, intracortical microelectrodes in a tetraplegic patient, J Neural Eng 18(4) (2021).

[27] X. Chen, F. Wang, R. Kooijmans, P.C. Klink, C. Boehler, M. Asplund, P.R. Roelfsema, Chronic stability of a neuroprosthesis comprising multiple adjacent Utah arrays in monkeys, J Neural Eng 20(3) (2023).

[28] J.P. Seymour, D.R. Kipke, Neural probe design for reduced tissue encapsulation in CNS, Biomaterials 28(25) (2007) 3594–3607.

[29] T.D. Kozai, D.R. Kipke, Insertion shuttle with carboxyl terminated self-assembled monolayer coatings for implanting flexible polymer neural probes in the brain, J Neurosci Methods 184(2) (2009) 199–205.

[30] J.P. Seymour, N.B. Langhals, D.J. Anderson, D.R. Kipke, Novel multi-sided, microelectrode arrays for implantable neural applications, Biomedical microdevices 13(3) (2011) 441–451.

[31] T.D.Y. Kozai, N.B. Langhals, P.R. Patel, X. Deng, H. Zhang, K.L. Smith, J. Lahann, N.A. Kotov, D.R. Kipke, Ultrasmall implantable composite microelectrodes with bioactive surfaces for chronic neural interfaces, Nat Mater 11(12) (2012) 1065–73.

[32] P. Gilgunn, R. Khilwani, T. Kozai, D. Weber, X. Cui, G. Erdos, O. Ozdoganlar, G. Fedder, An ultra-compliant, scalable neural probe with molded biodissolvable delivery vehicle, 2012 IEEE 25th international conference on micro electro mechanical systems (MEMS), IEEE, 2012, pp. 56-59.

[33] T.D.Y. Kozai, K. Catt, Z. Du, K. Na, O. Srivannavit, R.-U.M. Haque, J. Seymour, K.D. Wise, E. Yoon, X.T. Cui, Chronic In Vivo Evaluation of PEDOT/CNT for Stable Neural Recordings, IEEE transactions on bio-medical engineering 63(1) (2016) 111–119.

[34] S.M. Wellman, J.R. Eles, K.A. Ludwig, J.P. Seymour, N.J. Michelson, W.E. McFadden, A.L. Vazquez, T.D. Kozai, A Materials Roadmap to Functional Neural Interface Design, Advanced Functional Materials 28(12) (2018) 201701269.

[35] L. Luan, J.T. Robinson, B. Aazhang, T. Chi, K. Yang, X. Li, H. Rathore, A. Singer, S. Yellapantula, Y. Fan, Recent advances in electrical neural interface engineering: minimal invasiveness, longevity, and scalability, Neuron 108(2) (2020) 302–321.

[36] H. Steins, M. Mierzejewski, L. Brauns, A. Stumpf, A. Kohler, G. Heusel, A. Corna, T. Herrmann, P.D. Jones, G. Zeck, R. von Metzen, T. Stieglitz, A flexible protruding microelectrode array for neural interfacing in bioelectronic medicine, Microsystems & Nanoengineering 8(1) (2022) 131.

[37] I.N. McNamara, S.M. Wellman, L. Li, J.R. Eles, S. Savya, H.S. Sohal, M.R. Angle, T.D. Kozai, Electrode sharpness and insertion speed reduce tissue damage near high-density penetrating arrays, Journal of Neural Engineering 21(2) (2024) 026030.

[38] E. Musk, Neuralink, An Integrated Brain-Machine Interface Platform With Thousands of Channels, J Med Internet Res 21(10) (2019) e16194.

[39] V.S. Polikov, P.A. Tresco, W.M. Reichert, Response of brain tissue to chronically implanted neural electrodes, J Neurosci Methods 148(1) (2005) 1–18.

[40] K.C. Spencer, J.C. Sy, R. Falcon-Banchs, M.J. Cima, A three dimensional in vitro glial scar model to investigate the local strain effects from micromotion around neural implants, Lab Chip 17(5) (2017) 795–804.

[41] J.W. Salatino, K.A. Ludwig, T.D.Y. Kozai, E.K. Purcell, Glial responses to implanted electrodes in the brain, Nat Biomed Eng 1(11) (2017) 862–877.

[42] J. Duncan, A. Sridharan, S.S. Kumar, D. Iradukunda, J. Muthuswamy, Biomechanical micromotion at the neural interface modulates intracellular membrane potentialsin vivo, J Neural Eng 18(4) (2021).

[43] A. Trotier, E. Bagnoli, T. Walski, J. Evers, E. Pugliese, M. Lowery, M. Kilcoyne, U. Fitzgerald, M. Biggs, Micromotion Derived Fluid Shear Stress Mediates Peri-Electrode Gliosis through Mechanosensitive Ion Channels, Adv Sci (Weinh) 10(27) (2023) e2301352.

[44] R. Biran, D.C. Martin, P.A. Tresco, The brain tissue response to implanted silicon microelectrode arrays is increased when the device is tethered to the skull, J Biomed Mater Res A 82(1) (2007) 169–78.

[45] K. Chen, A. Forrest, G. Gonzalez Burgos, T.D.Y. Kozai, Neuronal functional connectivity is impaired in a layer dependent manner near chronically implanted intracortical microelectrodes in C57BL6 wildtype mice, Journal of Neural Engineering (2024).

[46] M. Dubaniewicz, J.R. Eles, S. Lam, S. Song, F. Cambi, D. Sun, S.M. Wellman, T.D. Kozai, Inhibition of Na+/H+ exchanger modulates microglial activation and scar formation following microelectrode implantation, Journal of Neural Engineering 18(4) (2021) 045001.

[47] J.R. Eles, A.L. Vazquez, N.R. Snyder, C.F. Lagenaur, M.C. Murphy, T.D.Y. Kozai, X.T. Cui, Neuroadhesive L1 coating attenuates acute microglial attachment to neural electrodes as revealed by live two-photon microscopy, Biomaterials 113 (2017) 279–292.

[48] T.D. Kozai, A.L. Vazquez, C.L. Weaver, S.G. Kim, X.T. Cui, In vivo two-photon microscopy reveals immediate microglial reaction to implantation of microelectrode through extension of processes, J Neural Eng 9(6) (2012) 066001.

[49] S.P. Savya, F. Li, S. Lam, S.M. Wellman, K.C. Stieger, K. Chen, J.R. Eles, T.D. Kozai, In vivo spatiotemporal dynamics of astrocyte reactivity following neural electrode implantation, Biomaterials 289 (2022) 121784.

[50] F. Li, J. Gallego, N.N. Tirko, J. Greaser, D. Bashe, R. Patel, E. Shaker, G.E. Van Valkenburg, A.S. Alsubhi, S. Wellman, Low-intensity pulsed ultrasound stimulation (LIPUS) modulates microglial activation following intracortical microelectrode implantation, Nature communications 15(1) (2024) 5512.

[51] T.D.Y. Kozai, A.S. Jaquins-gerstl, A.L. Vazquez, A.C. Michael, X.T. Cui, Dexamethasone retrodialysis attenuates microglial response to implanted probes in vivo, Biomaterials 87 (2016) 157–169.

[52] N.J. Michelson, A.L. Vazquez, J.R. Eles, J.W. Salatino, E.K. Purcell, J.J. Williams, X.T. Cui, T.D.Y. Kozai, Multi-scale, multi-modal analysis uncovers complex relationship at the brain tissue-implant neural interface: New Emphasis on the Biological Interface, Journal of Neural Engineering 15(033001) (2018).

[53] B. Coste, J. Mathur, M. Schmidt, T.J. Earley, S. Ranade, M.J. Petrus, A.E. Dubin, A. Patapoutian, Piezo1 and Piezo2 are essential components of distinct mechanically activated cation channels, Science 330(6000) (2010) 55-60.

[54] H. Liu, J. Hu, Q. Zheng, X. Feng, F. Zhan, X. Wang, G. Xu, F. Hua, Piezo1 Channels as Force Sensors in Mechanical Force-Related Chronic Inflammation, Front Immunol 13 (2022) 816149.

[55] P. Moshayedi, G. Ng, J.C. Kwok, G.S. Yeo, C.E. Bryant, J.W. Fawcett, K. Franze, J. Guck, The relationship between glial cell mechanosensitivity and foreign body reactions in the central nervous system, Biomaterials 35(13) (2014) 3919–25.

[56] M.S. Fee, Active stabilization of electrodes for intracellular recording in awake behaving animals, Neuron 27(3) (2000) 461–8.

[57] A. Gilletti, J. Muthuswamy, Brain micromotion around implants in the rodent somatosensory cortex, J Neural Eng 3(3) (2006) 189–95.

[58] H. Jantti, V. Sitnikova, Y. Ishchenko, A. Shakirzyanova, L. Giudice, I.F. Ugidos, M. Gomez-Budia, N. Korvenlaita, S. Ohtonen, I. Belaya, F. Fazaludeen, N. Mikhailov, M. Gotkiewicz, K. Ketola, S. Lehtonen, J. Koistinaho, K.M. Kanninen, D. Hernandez, A. Pebay, R. Giugno, P. Korhonen, R. Giniatullin, T. Malm, Microglial amyloid beta clearance is driven by PIEZO1 channels, J Neuroinflammation 19(1) (2022) 147.

[59] J. Hu, Q. Chen, H. Zhu, L. Hou, W. Liu, Q. Yang, H. Shen, G. Chai, B. Zhang, S. Chen, Z. Cai, C. Wu, F. Hong, H. Li, S. Chen, N. Xiao, Z.X. Wang, X. Zhang, B. Wang, L. Zhang, W. Mo, Microglial Piezo1 senses Abeta fibril stiffness to restrict Alzheimer’s disease, Neuron 111(1) (2023) 15–29 e8.

[60] G. Hong, C.M. Lieber, Novel electrode technologies for neural recordings, Nat Rev Neurosci 20(6) (2019) 330–345.

[61] J.P. Harris, J.R. Capadona, R.H. Miller, B.C. Healy, K. Shanmuganathan, S.J. Rowan, C. Weder, D.J. Tyler, Mechanically adaptive intracortical implants improve the proximity of neuronal cell bodies, J Neural Eng 8(6) (2011) 066011.

[62] C.L. Kolarcik, S.D. Luebben, S.A. Sapp, J. Hanner, N. Snyder, T.D. Kozai, E. Chang, J.A. Nabity, S.T. Nabity, C.F. Lagenaur, X.T. Cui, Elastomeric and soft conducting microwires for implantable neural interfaces, Soft Matter 11(24) (2015) 4847–61.

[63] A. Sridharan, J.K. Nguyen, J.R. Capadona, J. Muthuswamy, Compliant intracortical implants reduce strains and strain rates in brain tissue in vivo, J Neural Eng 12(3) (2015) 036002.

[64] R. Khilwani, P.J. Gilgunn, T.D. Kozai, X.C. Ong, E. Korkmaz, P.K. Gunalan, X.T. Cui, G.K. Fedder, O.B. Ozdoganlar, Ultra- miniature ultra-compliant neural probes with dissolvable delivery needles: design, fabrication and characterization, Biomed Microdevices 18(6) (2016) 97.

[65] Z.J. Du, C.L. Kolarcik, T.D.Y. Kozai, S.D. Luebben, S.A. Sapp, X.S. Zheng, J.A. Nabity, X.T. Cui, Ultrasoft microwire neural electrodes improve chronic tissue integration, Acta Biomater 53 (2017) 46–58.

[66] H.C. Lee, F. Ejserholm, J. Gaire, S. Currlin, J. Schouenborg, L. Wallman, M. Bengtsson, K. Park, K.J. Otto, Histological evaluation of flexible neural implants; flexibility limit for reducing the tissue response?, J Neural Eng 14(3) (2017) 036026.

[67] A.M. Stiller, J.O. Usoro, J. Lawson, B. Araya, M.A. Gonzalez-Gonzalez, V.R. Danda, W.E. Voit, B.J. Black, J.J. Pancrazio, Mechanically Robust, Softening Shape Memory Polymer Probes for Intracortical Recording, Micromachines (Basel) 11(6) (2020).

[68] N. Sharafkhani, J.M. Long, S.D. Adams, A.Z. Kouzani, A self-stiffening compliant intracortical microprobe, Biomed Microdevices 26(1) (2024) 17.

[69] A.G. Solis, P. Bielecki, H.R. Steach, L. Sharma, C.C.D. Harman, S. Yun, M.R. de Zoete, J.N. Warnock, S.D.F. To, A.G. York, M. Mack, M.A. Schwartz, C.S.D. Cruz, N.W. Palm, R. Jackson, R.A. Flavell, Author Correction: Mechanosensation of cyclical force by PIEZO1 is essential for innate immunity, Nature 575(7784) (2019) E7.

[70] H. Atcha, A. Jairaman, J.R. Holt, V.S. Meli, R.R. Nagalla, P.K. Veerasubramanian, K.T. Brumm, H.E. Lim, S. Othy, M.D. Cahalan, M.M. Pathak, W.F. Liu, Mechanically activated ion channel Piezo1 modulates macrophage polarization and stiffness sensing, Nat Commun 12(1) (2021) 3256.

[71] R.G. Scheraga, S. Abraham, K.A. Niese, B.D. Southern, L.M. Grove, R.D. Hite, C. McDonald, T.A. Hamilton, M.A. Olman, TRPV4 Mechanosensitive Ion Channel Regulates Lipopolysaccharide-Stimulated Macrophage Phagocytosis, J Immunol 196(1) (2016) 428–36.

[72] D. Yu, A. Ahmed, J. Jayasi, A. Womac, O. Sally, C. Bae, Inflammation condition sensitizes Piezo1 mechanosensitive channel in mouse cerebellum astrocyte, Front Cell Neurosci 17 (2023) 1200946.

[73] S. Chi, Y. Cui, H. Wang, J. Jiang, T. Zhang, S. Sun, Z. Zhou, Y. Zhong, B. Xiao, Astrocytic Piezo1-mediated mechanotransduction determines adult neurogenesis and cognitive functions, Neuron 110(18) (2022) 2984–2999 e8.

[74] M. Velasco-Estevez, K.K.E. Gadalla, N. Linan-Barba, S. Cobb, K.K. Dev, G.K. Sheridan, Inhibition of Piezo1 attenuates demyelination in the central nervous system, Glia 68(2) (2020) 356–375.

[75] B.T. Jacques-Fricke, Y. Seow, P.A. Gottlieb, F. Sachs, T.M. Gomez, Ca2+ influx through mechanosensitive channels inhibits neurite outgrowth in opposition to other influx pathways and release from intracellular stores, J Neurosci 26(21) (2006) 5656–64.

[76] J.R. Eles, A.L. Vazquez, T.D.Y. Kozai, X.T. Cui, In vivo imaging of neuronal calcium during electrode implantation: Spatial and temporal mapping of damage and recovery, Biomaterials 174 (2018) 79–94.

[77] C.S. Bjornsson, S.J. Oh, Y.A. Al-Kofahi, Y.J. Lim, K.L. Smith, J.N. Turner, S. De, B. Roysam, W. Shain, S.J. Kim, Effects of insertion conditions on tissue strain and vascular damage during neuroprosthetic device insertion, J Neural Eng 3(3) (2006) 196–207.

[78] M.D. Johnson, O.E. Kao, D.R. Kipke, Spatiotemporal pH dynamics following insertion of neural microelectrode arrays, Journal of neuroscience methods 160(2) (2007) 276–287.

[79] T.D. Kozai, T.C. Marzullo, F. Hooi, N.B. Langhals, A.K. Majewska, E.B. Brown, D.R. Kipke, Reduction of neurovascular damage resulting from microelectrode insertion into the cerebral cortex using in vivo two-photon mapping, J Neural Eng 7(4) (2010) 046011.

[80] T. Escamilla-Mackert, N.B. Langhals, T.D. Kozai, D.R. Kipke, Insertion of a three dimensional silicon microelectrode assembly through a thick meningeal membrane, Annu Int Conf IEEE Eng Med Biol Soc 2009 (2009) 1616-8.

[81] K. Sahasrabuddhe, A.A. Khan, A.P. Singh, T.M. Stern, Y. Ng, A. Tadić, P. Orel, C. LaReau, D. Pouzzner, K. Nishimura, K.M. Boergens, S. Shivakumar, M.S. Hopper, B. Kerr, M.-E.S. Hanna, R.J. Edgington, I. McNamara, D. Fell, P. Gao, A. Babaie-Fishani, S. Veijalainen, A.V. Klekachev, A.M. Stuckey, B. Luyssaert, T.D.Y. Kozai, C. Xie, V. Gilja, B. Dierickx, Y. Kong, M. Straka, H.S. Sohal, M.R. Angle, The Argo: a high channel count recording system for neural recording in vivo, Journal of Neural Engineering 18(1) (2021) 015002.

[82] X. Chen, A. Morales-Gregorio, J. Sprenger, A. Kleinjohann, S. Sridhar, S.J. van Albada, S. Grün, P.R. Roelfsema, 1024- channel electrophysiological recordings in macaque V1 and V4 during resting state, Scientific Data 9(1) (2022) 77.

[83] A. Obaid, Y.-W. Wu, M. Hanna, O. Jáidar, W. Nix, J. Ding, N. Melosh, Ultra-sensitive measurement of brain penetration mechanics and blood vessel rupture with microscale probes, bioRxiv (2020) 2020.09. 21.306498.

[84] H. Lee, R.V. Bellamkonda, W. Sun, M.E. Levenston, Biomechanical analysis of silicon microelectrode-induced strain in the brain, J Neural Eng 2(4) (2005) 81–9.

[85] J. Subbaroyan, D.C. Martin, D.R. Kipke, A finite-element model of the mechanical effects of implantable microelectrodes in the cerebral cortex, J Neural Eng 2(4) (2005) 103–13.

[86] T.D. Kozai, K. Catt, X. Li, Z.V. Gugel, V.T. Olafsson, A.L. Vazquez, X.T. Cui, Mechanical failure modes of chronically implanted planar silicon-based neural probes for laminar recording, Biomaterials 37 (2015) 25–39.

[87] M. Polanco, S. Bawab, H. Yoon, Computational Assessment of Neural Probe and Brain Tissue Interface under Transient Motion, Biosensors (Basel) 6(2) (2016) 27.

[88] W. Zhang, J. Tang, Z. Li, Y. Ma, A novel neural electrode with micro-motion-attenuation capability based on compliant mechanisms-physical design concepts and evaluations, Med Biol Eng Comput 56(10) (2018) 1911–1923.

[89] A. Al Abed, J. Amatoury, M. Khraiche, Finite Element Modeling of Magnitude and Location of Brain Micromotion Induced Strain for Intracortical Implants, Front Neurosci 15 (2021) 727715.

[90] K. Kim, C. Sung, J. Lee, J. Won, W. Jeon, S. Seo, K. Yoon, S. Park, Computational and Histological Analyses for Investigating Mechanical Interaction of Thermally Drawn Fiber Implants with Brain Tissue, Micromachines (Basel) 12(4) (2021).

[91] N.H. Hosseini, R. Hoffmann, S. Kisban, T. Stieglitz, O. Paul, P. Ruther, Comparative study on the insertion behavior of cerebral microprobes, Annu Int Conf IEEE Eng Med Biol Soc 2007 (2007) 4711–4.

[92] S. Budday, G. Sommer, C. Birkl, C. Langkammer, J. Haybaeck, J. Kohnert, M. Bauer, F. Paulsen, P. Steinmann, E. Kuhl, G.A. Holzapfel, Mechanical characterization of human brain tissue, Acta Biomater 48 (2017) 319–340.

[93] D.A. Henze, Z. Borhegyi, J. Csicsvari, A. Mamiya, K.D. Harris, G. Buzsaki, Intracellular features predicted by extracellular recordings in the hippocampus in vivo, J Neurophysiol 84(1) (2000) 390–400.

[94] G. Buzsaki, Large-scale recording of neuronal ensembles, Nat Neurosci 7(5) (2004) 446–51.

[95] S.N. Flesher, J.E. Downey, J.M. Weiss, C.L. Hughes, A.J. Herrera, E.C. Tyler-Kabara, M.L. Boninger, J.L. Collinger, R.A. Gaunt, A brain-computer interface that evokes tactile sensations improves robotic arm control, Science 372(6544) (2021) 831-836.

[96] B.M. Dekleva, J.M. Weiss, M.L. Boninger, J.L. Collinger, Generalizable cursor click decoding using grasp-related neural transients, J Neural Eng 18(4) (2021).

[97] J.E. Downey, H.R. Schone, S.T. Foldes, C. Greenspon, F. Liu, C. Verbaarschot, D. Biro, D. Satzer, C.H. Moon, B.A. Coffman, V. Youssofzadeh, D. Fields, T.G. Hobbs, E. Okorokova, E.C. Tyler-Kabara, P.C. Warnke, J. Gonzalez-Martinez, N.G. Hatsopoulos, S.J. Bensmaia, M.L. Boninger, R.A. Gaunt, J.L. Collinger, A Roadmap for Implanting Electrode Arrays to Evoke Tactile Sensations Through Intracortical Stimulation, Hum Brain Mapp 45(18) (2024) e70118.

[98] N.G. Kunigk, H.R. Schone, C. Gontier, W. Hockeimer, A.F. Tortolani, N.G. Hatsopoulos, J.E. Downey, S.M. Chase, M.L. Boninger, B.D. Dekleva, J.L. Collinger, Motor somatotopy impacts imagery strategy success in human intracortical brain-computer interfaces, J Neural Eng 22(2) (2025).

[99] X. Chen, F. Wang, E. Fernandez, P.R. Roelfsema, Shape perception via a high-channel-count neuroprosthesis in monkey visual cortex, Science 370(6521) (2020) 1191-1196.

[100] H. Super, P.R. Roelfsema, Chronic multiunit recordings in behaving animals: advantages and limitations, Prog Brain Res 147 (2005) 263–82.

[101] R. Biran, D.C. Martin, P.A. Tresco, Neuronal cell loss accompanies the brain tissue response to chronically implanted silicon microelectrode arrays, Exp Neurol 195(1) (2005) 115–26.

[102] G.C. McConnell, H.D. Rees, A.I. Levey, C.A. Gutekunst, R.E. Gross, R.V. Bellamkonda, Implanted neural electrodes cause chronic, local inflammation that is correlated with local neurodegeneration, J Neural Eng 6(5) (2009) 56003.

[103] S.M. Wellman, L. Li, Y. Yaxiaer, I. McNamara, T.D.Y. Kozai, Revealing Spatial and Temporal Patterns of Cell Death, Glial Proliferation, and Blood-Brain Barrier Dysfunction Around Implanted Intracortical Neural Interfaces, Front Neurosci 13 (2019) 493.

[104] L. Bollmann, D.E. Koser, R. Shahapure, H.O. Gautier, G.A. Holzapfel, G. Scarcelli, M.C. Gather, E. Ulbricht, K. Franze, Microglia mechanics: immune activation alters traction forces and durotaxis, Front Cell Neurosci 9 (2015) 363.

[105] J. Liu, Y. Yang, Y. Liu, Piezo1 plays a role in optic nerve head astrocyte reactivity, Exp Eye Res 204 (2021) 108445.

[106] N.R. Patel, M. Bole, C. Chen, C.C. Hardin, A.T. Kho, J. Mih, L. Deng, J. Butler, D. Tschumperlin, J.J. Fredberg, R. Krishnan, H. Koziel, Cell elasticity determines macrophage function, PLoS One 7(9) (2012) e41024.

[107] M.B. Christensen, H.A. Wark, D.T. Hutchinson, A histological analysis of human median and ulnar nerves following implantation of Utah slanted electrode arrays, Biomaterials 77 (2016) 235–42.

[108] K. Woeppel, C. Hughes, A.J. Herrera, J.R. Eles, E.C. Tyler-Kabara, R.A. Gaunt, J.L. Collinger, X.T. Cui, Explant Analysis of Utah Electrode Arrays Implanted in Human Cortex for Brain-Computer-Interfaces, Front Bioeng Biotechnol 9 (2021) 759711.

[109] P.R. Patel, E.J. Welle, J.G. Letner, H. Shen, A.J. Bullard, C.M. Caldwell, A. Vega-Medina, J.M. Richie, H.E. Thayer, P.G. Patil, D. Cai, C.A. Chestek, Utah array characterization and histological analysis of a multi-year implant in non-human primate motor and sensory cortices, J Neural Eng 20(1) (2023).

[110] T.D.Y. Kozai, A. Jaquins-Gerstl, A.L. Vazquez, A.C. Michael, X.T. Cui, Brain Tissue Responses to Neural Implants Impact Signal Sensitivity and Intervention Strategies, ACS Chemical Neuroscience 6(1) (2015) 48–67.

[111] S.P. Burns, D. Xing, R.M. Shapley, Comparisons of the dynamics of local field potential and multiunit activity signals in macaque visual cortex, J Neurosci 30(41) (2010) 13739–49.

[112] Y.S. Choi, M.A. Koenig, X. Jia, N.V. Thakor, Quantifying time-varying multiunit neural activity using entropy based measures, IEEE Trans Biomed Eng 57(11) (2010).

[113] E. Cetinkaya, E.J. Lang, M. Sahin, Sensorimotor content of multi-unit activity recorded in the paramedian lobule of the cerebellum using carbon fiber microelectrode arrays, Front Neurosci 18 (2024) 1232653.

[114] K. Torab, T.S. Davis, D.J. Warren, P.A. House, R.A. Normann, B. Greger, Multiple factors may influence the performance of a visual prosthesis based on intracortical microstimulation: nonhuman primate behavioural experimentation, J Neural Eng 8(3) (2011) 035001.

[115] S.R. Kane, S.F. Cogan, J. Ehrlich, T.D. Plante, D.B. McCreery, P.R. Troyk, Electrical performance of penetrating microelectrodes chronically implanted in cat cortex, IEEE Trans Biomed Eng 60(8) (2013) 2153–60.

[116] T.D. Kozai, X. Li, L.M. Bodily, E.M. Caparosa, G.A. Zenonos, D.L. Carlisle, R.M. Friedlander, X.T. Cui, Effects of caspase-1 knockout on chronic neural recording quality and longevity: insight into cellular and molecular mechanisms of the reactive tissue response, Biomaterials 35(36) (2014) 9620–34.

[117] J.P. Vande Geest, B.R. Simon, P.H. Rigby, T.P. Newberg, Coupled porohyperelastic mass transport (PHEXPT) finite element models for soft tissues using ABAQUS, J Biomech Eng 133(4) (2011) 044502.

[118] J.T. Keyes, B.R. Simon, J.P. Vande Geest, Location-dependent coronary artery diffusive and convective mass transport properties of a lipophilic drug surrogate measured using nonlinear microscopy, Pharm Res 30(4) (2013) 1147–60.

[119] J.T. Keyes, D.R. Lockwood, B.R. Simon, J.P. Vande Geest, Deformationally dependent fluid transport properties of porcine coronary arteries based on location in the coronary vasculature, J Mech Behav Biomed Mater 17 (2013) 296–306.

[120] A. Ayyalasomayajula, R.I. Park, B.R. Simon, J.P. Vande Geest, A porohyperelastic finite element model of the eye: the influence of stiffness and permeability on intraocular pressure and optic nerve head biomechanics, Comput Methods Biomech Biomed Engin 19(6) (2016) 591–602.

[121] M.H. Armstrong, A. Buganza Tepole, E. Kuhl, B.R. Simon, J.P. Vande Geest, A Finite Element Model for Mixed Porohyperelasticity with Transport, Swelling, and Growth, PLoS One 11(4) (2016) e0152806.

[122] A. Prasad, Q.-S. Xue, R. Dieme, V. Sankar, R.C. Mayrand, T. Nishida, W.J. Streit, J.C. Sanchez, Abiotic-biotic characterization of Pt/Ir microelectrode arrays in chronic implants, Frontiers in neuroengineering 7 (2014) 2.

[123] S. Kim, R.A. Normann, R. Harrison, F. Solzbacher, Preliminary study of the thermal impact of a microelectrode array implanted in the brain, Conf Proc IEEE Eng Med Biol Soc 2006 (2006) 2986–9.

[124] S. Kim, P. Tathireddy, R.A. Normann, F. Solzbacher, Thermal impact of an active 3-D microelectrode array implanted in the brain, IEEE Trans Neural Syst Rehabil Eng 15(4) (2007) 493–501.

[125] L. Luan, X. Wei, Z. Zhao, J.J. Siegel, O. Potnis, C.A. Tuppen, S. Lin, S. Kazmi, R.A. Fowler, S. Holloway, A.K. Dunn, R.A. Chitwood, C. Xie, Ultraflexible nanoelectronic probes form reliable, glial scar-free neural integration, Sci Adv 3(2) (2017) e1601966.

[126] J. Shi, Y. Fang, Flexible and Implantable Microelectrodes for Chronically Stable Neural Interfaces, Adv Mater 31(45) (2019) e1804895.

[127] K.C. Stocking, A.L. Vazquez, T.D.Y. Kozai, Intracortical Neural Stimulation With Untethered, Ultrasmall Carbon Fiber Electrodes Mediated by the Photoelectric Effect, IEEE Trans Biomed Eng 66(8) (2019) 2402–2412.

[128] A. Golabchi, K.M. Woeppel, X. Li, C.F. Lagenaur, X.T. Cui, Neuroadhesive protein coating improves the chronic performance of neuroelectronics in mouse brain, Biosens Bioelectron 155 (2020) 112096.

[129] N.S. Witham, C.F. Reiche, T. Odell, K. Barth, C.H. Chiang, C. Wang, A. Dubey, K. Wingel, S. Devore, D. Friedman, B. Pesaran, J. Viventi, F. Solzbacher, Flexural bending to approximate cortical forces exerted by electrocorticography (ECoG) arrays, J Neural Eng 19(4) (2022).

[130] M.E. Urdaneta, N.G. Kunigk, F. Delgado, S.I. Fried, K.J. Otto, Layer-specific parameters of intracortical microstimulation of the somatosensory cortex, J Neural Eng 18(5) (2021).

[131] M.E. Urdaneta, N.G. Kunigk, J.D. Penaloza-Aponte, S. Currlin, I.G. Malone, S.I. Fried, K.J. Otto, Layer-dependent stability of intracortical recordings and neuronal cell loss, Front Neurosci 17 (2023) 1096097.

[132] J.P. Harris, A.E. Hess, S.J. Rowan, C. Weder, C.A. Zorman, D.J. Tyler, J.R. Capadona, In vivo deployment of mechanically adaptive nanocomposites for intracortical microelectrodes, J Neural Eng 8(4) (2011) 046010.

[133] P.J. Rousche, R.A. Normann, A method for pneumatically inserting an array of penetrating electrodes into cortical tissue, Ann Biomed Eng 20(4) (1992) 413–22.

[134] P.A. House, J.D. MacDonald, P.A. Tresco, R.A. Normann, Acute microelectrode array implantation into human neocortex: preliminary technique and histological considerations, Neurosurg Focus 20(5) (2006) E4.

[135] K.P. O’Sullivan, B. Coats, Coupled Eulerian-Lagrangian model prediction of neural tissue strain during microelectrode insertion, J Neural Eng (2024).

