## SupplementalFigures for "Finite element model predicts micromotion-induced strain profiles that correlate with the functional performance of Utah arrays in humans and non-human primates"

**Supplementary Figures**

Supplementary Figure 1. Convergence study to determine finite element modeling mesh size.
Supplementary Figure 2. Human motor array performance metrics
Supplementary Figure 3. Human somatosensory array performance metrics.
Supplementary Figure 4. Session-to-session variability shows some dependence on electrode location within the array.
Supplementary Figure 5. Non-human primate area V4 array performance metrics.
Supplementary Figure 6. Predicted maximal von Mises strains for 10x10 array geometry show similar trends as average strains.
Supplementary Figure 7. Predicted average von Mises strains for 10x6 and 8x8 array geometries are similar to 10x10 geometry.
Supplementary Figure 8. Average correlation with neighboring bandpass-filtered electrophysiological recordings.
Supplementary Figure 9. Spontaneous peak-to-peak voltage and SNR from implanted human somatosensory arrays show limited relationship with micromotion strains.

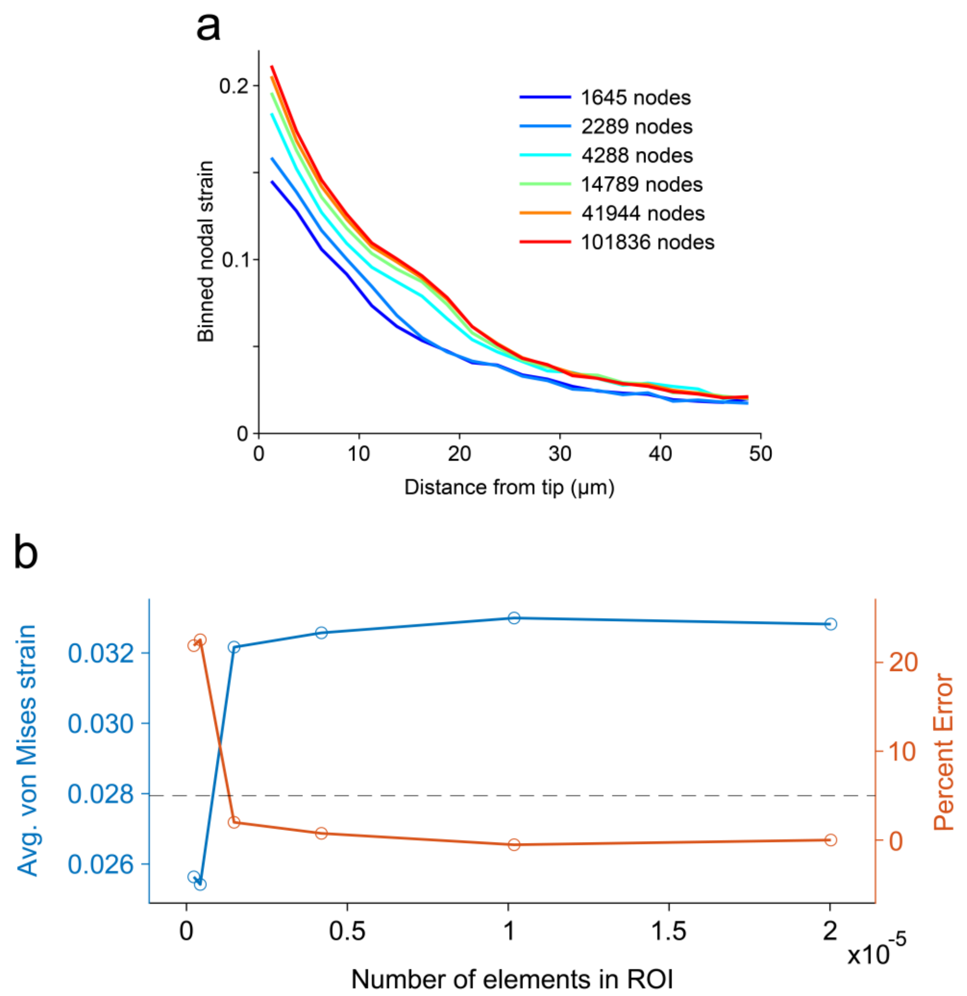


**Supplementary Figure 1. Convergence study to determine finite element modeling mesh size.** (a) Convergence study was conducted on a single electrode to reduce computation time. Element size surrounding the electrode tip was refined to achieve higher density meshes. Nodal strains were averaged in 2.5μm bins surrounding electrode tip. (b) Average von Mises strain, computed using the binned strains in (a), converges with increasing element number. The meshing parameters for the third mesh (with 4288 nodes) were selected as an optimal tradeoff of accuracy and computational time.

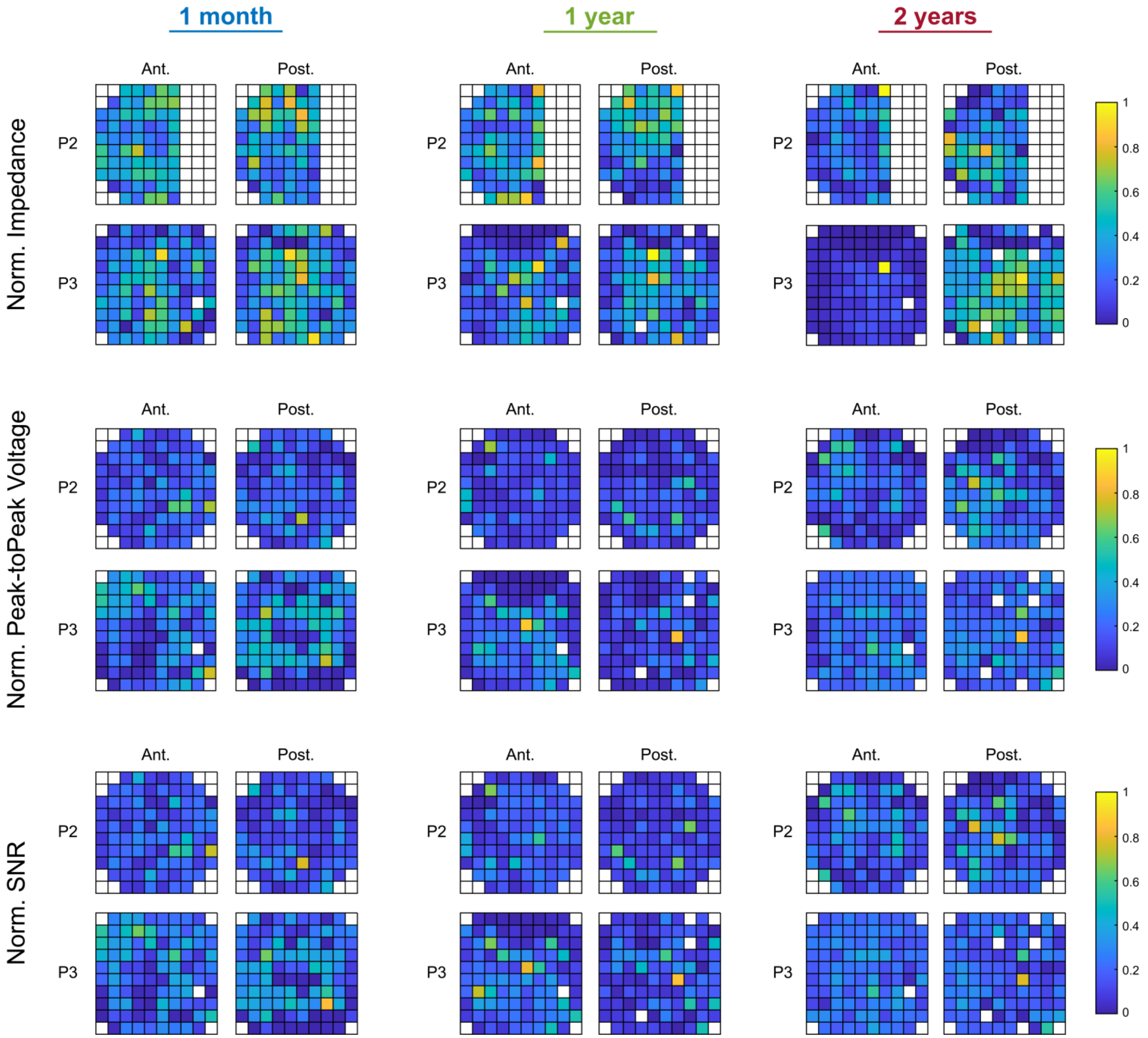


**Supplementary Figure 2. Human motor array performance metrics.** For each time period of interest (1-mo, 1-yr, and 2-yrs post-implantation) a total of five successive recording sessions were analyzed (N = 2 participants, n = 4 arrays). For each array, the performance measures were first averaged across the five recording sessions and then normalized within each array. Heatmaps show the normalized performance metrics for each array. Electrodes whited out were either not used for recordings or had 1kHz impedance greater than 1MΩ on every recording day. The large bank of electrodes missing impedance data from P2 were used for recording, but impedance measurements were not collected.

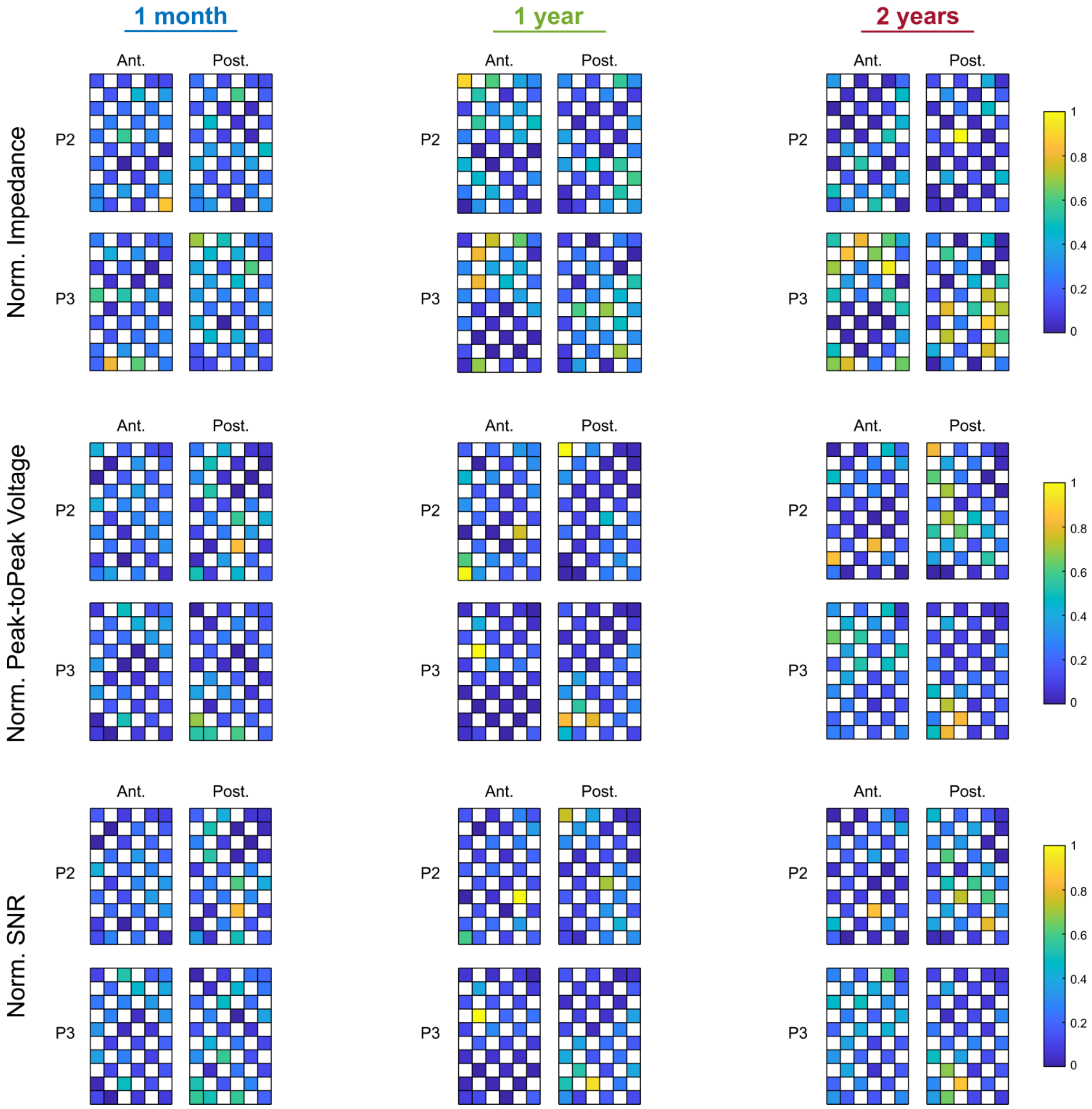


**Supplementary Figure 3. Human somatosensory array performance metrics.** For each time period of interest (1-mo, 1-yr, and 2-yrs post-implantation) a total of five successive recording sessions were analyzed (N = 2 participants, n = 4 arrays). For each array, the performance measures were first averaged across the five recording sessions and then normalized within each array. Heatmaps show the normalized performance metrics for each array. Electrodes whited out were not used for recordings.

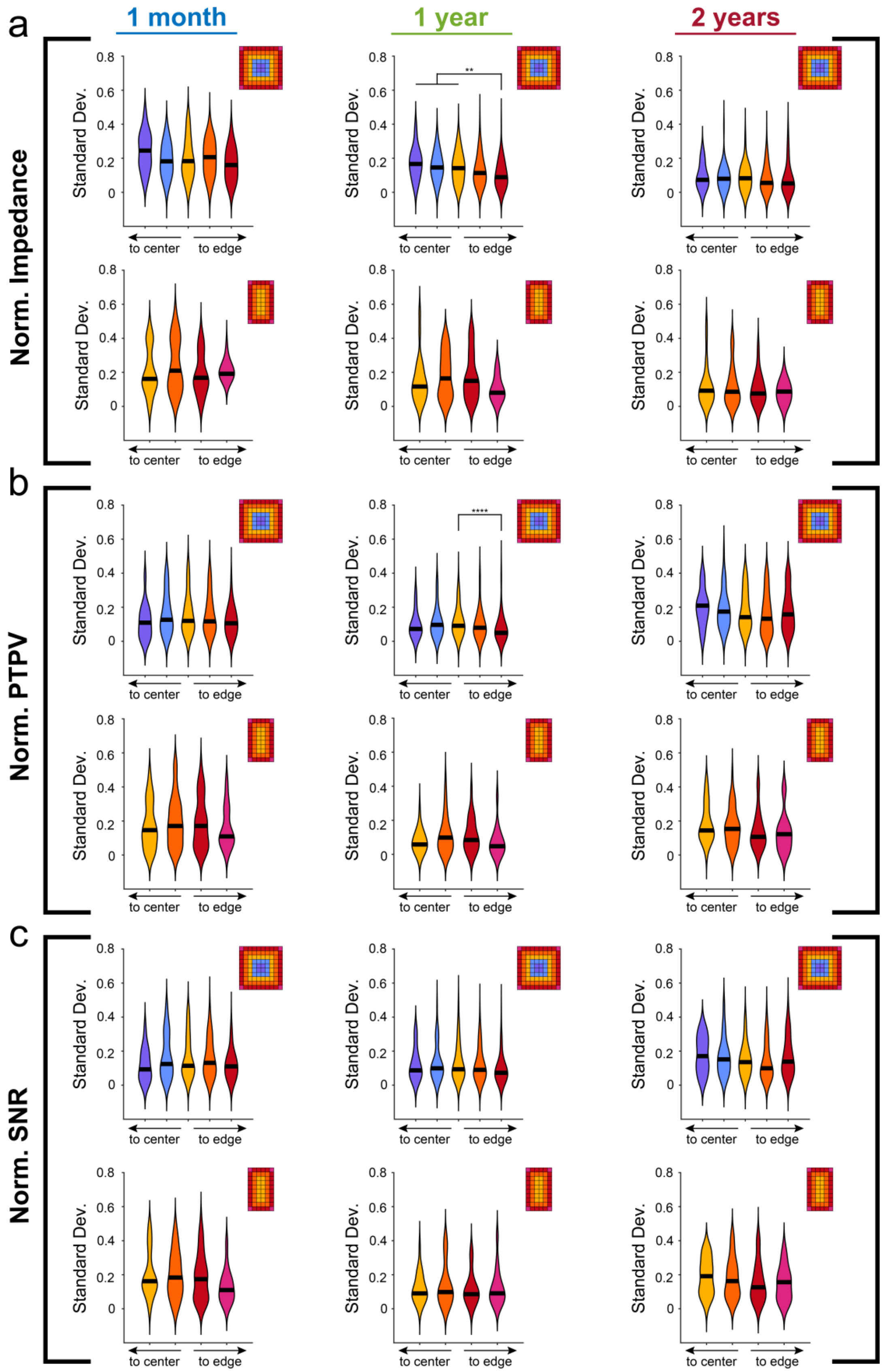


**Supplementary Figure 4. Session-to-session variability shows some dependence on electrode location within the array.** Violin plots represent standard deviations of (a) 1kHz impedance, (b) PTPV, and (c) SNR for each electrode across five successive recording sessions. For impedance and SNR metrics, electrodes whited-out are from electrodes whose 1kHz impedance was greater than 1MΩ. No significant differences in session-to-session variability between edge and interior electrodes were observed for the 10x6 somatosensory arrays (Kruskal-Wallis test with post-hoc Dunn’s test). For 10x10 motor arrays, variability in the impedance and PTPV of edge electrodes 1-yr post-implantation was significantly lower compared to interior electrodes (Kruskal-Wallis test with post-hoc Dunn’s test). ** $p<0.01$, *** $p<0.001$, **** $p<0.0001$.

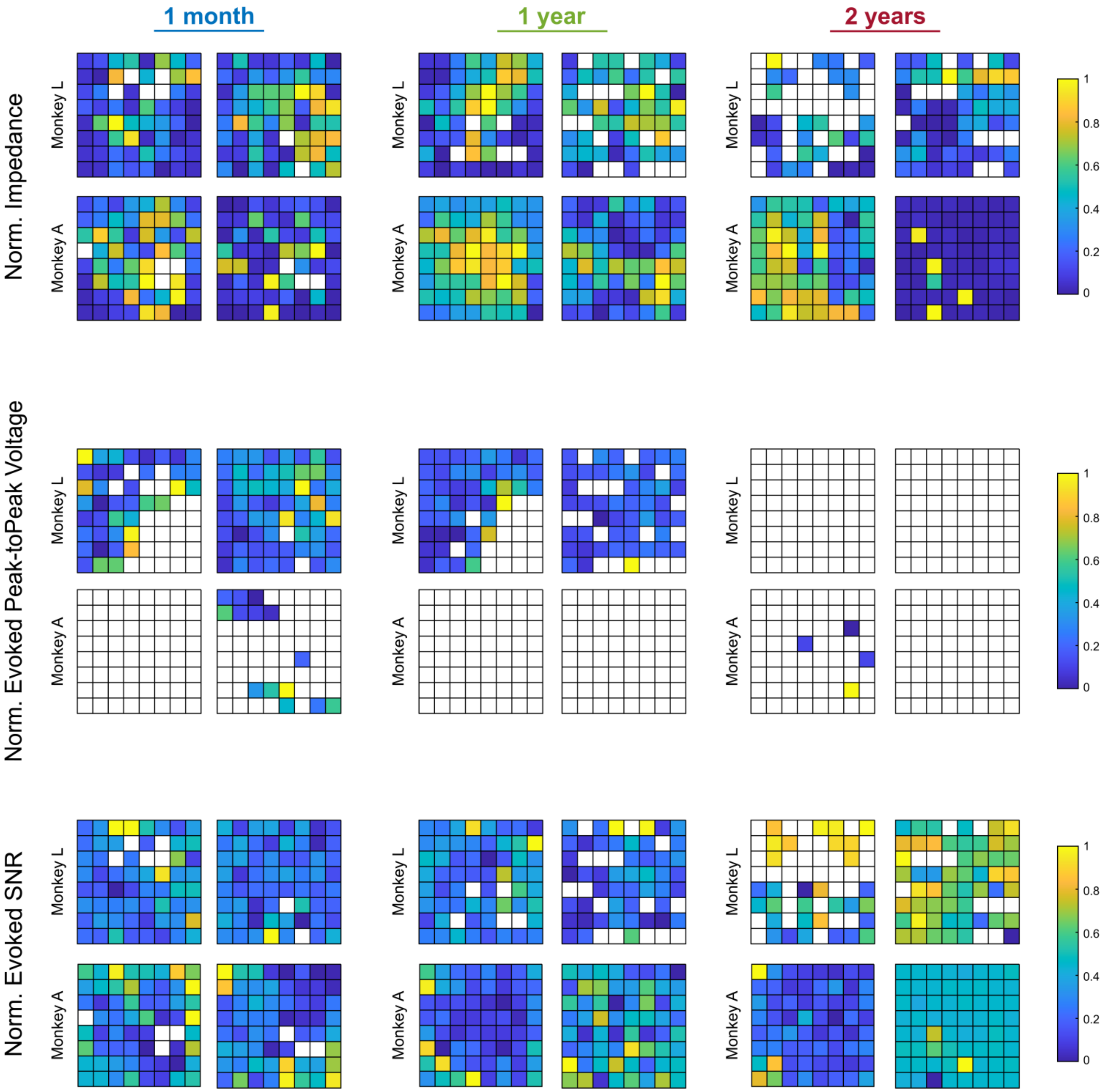


**Supplementary Figure 5. Non-human primate area V4 array performance metrics.** Heatmaps show normalized performance measures for each array (N = 2 animals, n = 4 arrays). Limited number of recording sessions prevented analysis for session-to-session variability. Electrodes whited out either had 1kHz impedance greater than 1MΩ or represent electrodes with very low PTPV amplitudes (less than 30µV) and were excluded from the PTPV analysis.

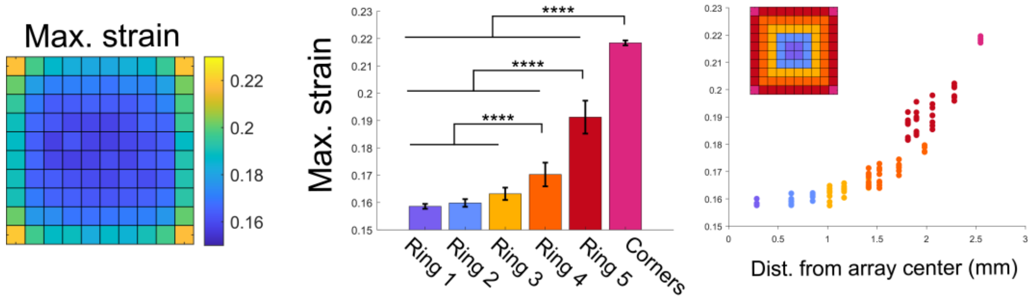


**Supplementary Figure 6. Predicted maximal von Mises strains for 10x10 array geometry show similar trends as average strains.** Heatmap of maximum von Mises strain occurring within 50µm of each electrode for 10x10 array (left). Maximum strains at corner, ring 5, and ring 4 electrodes were significantly different from all other groups ($p<0.0001$, one-way ANOVA with Tukey's post-hoc test). Maximum strains plotted as a function of distance from the center of the array show corner and ring 5 electrodes are distinctly grouped (right). Error bars are standard deviations. ** $p<0.01$, *** $p<0.001$, **** $p<0.0001$.

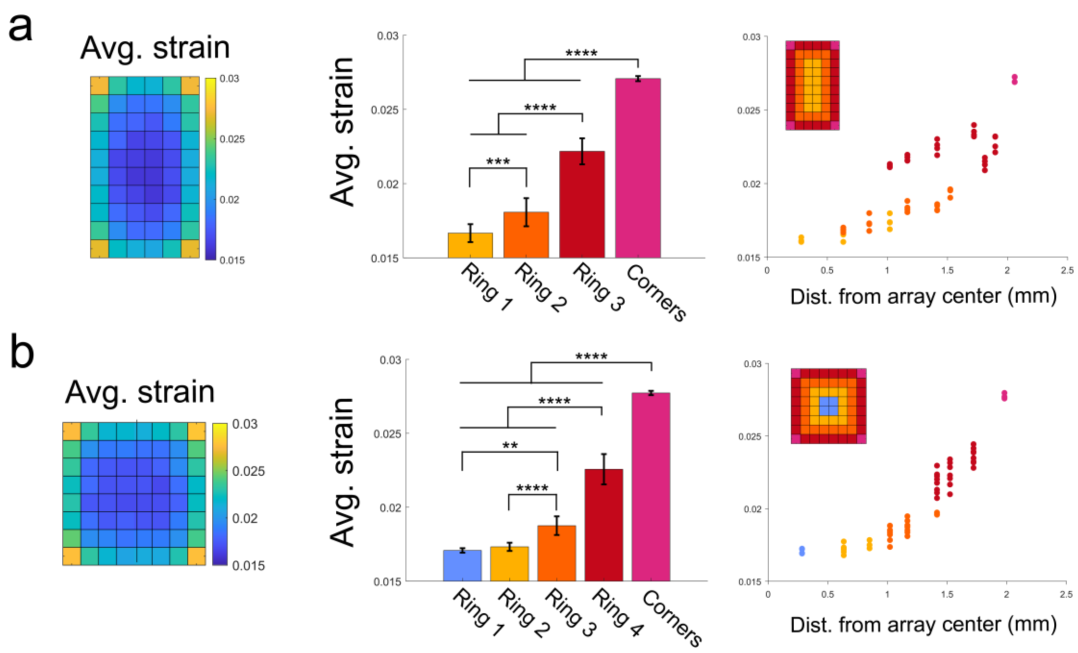


**Supplementary Figure 7. Predicted average von Mises strains for 10x6 and 8x8 array geometries are similar to 10x10 geometry**. (a) Heatmap of average von Mises strain occurring within 50µm of each electrode for 10x6 array (left). Average strains for each group were significantly different from all other groups (p<0.001, one-way ANOVA with Tukey's post-hoc test). Average strains plotted as a function of distance from the center of the array show corner and ring 3 electrodes are distinctly grouped (right). (b) Heatmap of average von Mises strain occurring within 50µm of each electrode for 8x8 array (left). Average strains at corner, ring 4, and ring 3 electrodes were significantly different from all other groups (p<0.01, one-way ANOVA with Tukey's post-hoc test). Average strains plotted as a function of distance from the center of the array show corner and ring 4 electrodes are distinctly grouped (right). Error bars are standard deviations. ** p<0.01, *** p<0.001, **** p<0.0001.

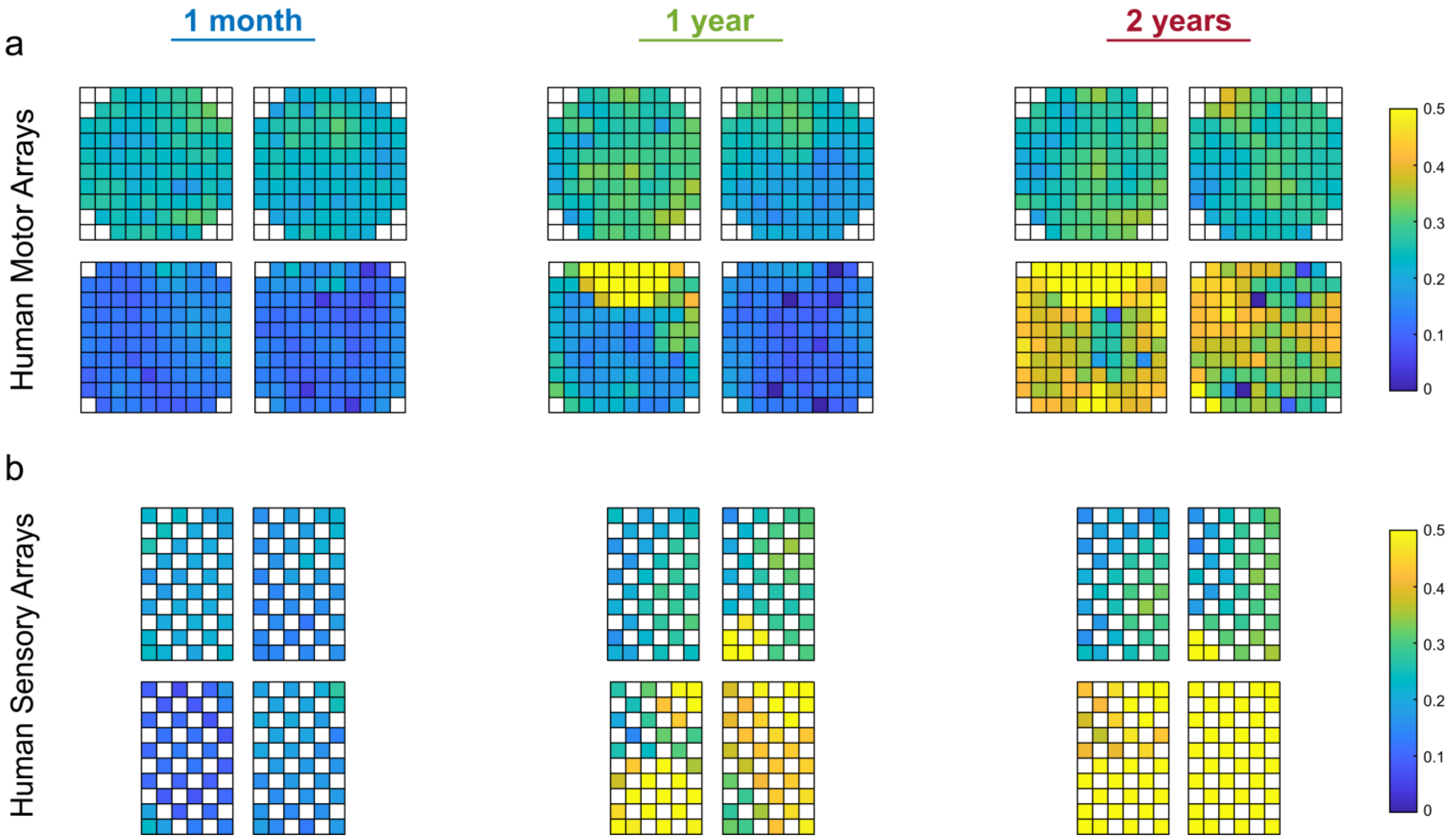


**Supplementary Figure 8. Average correlation with neighboring bandpass-filtered electrophysiological recordings.** Heatmaps for (a) 10x10 human motor arrays and (b) 10x6 human somatosensory arrays showing correlation between the bandpass-filtered electrophysiological recordings of neighboring electrodes for all arrays and studied timepoints. Briefly, for each electrode, its bandpass-filtered recording was correlated with 3-8 neighboring electrodes (depending on electrode location). The average of these 3-8 neighboring electrode correlations was then computed. This was repeated for five successive recording sessions for each time period. Heatmaps shown are averages across the five successive recording sessions and whose values are correlated with impedance in Figure 4.

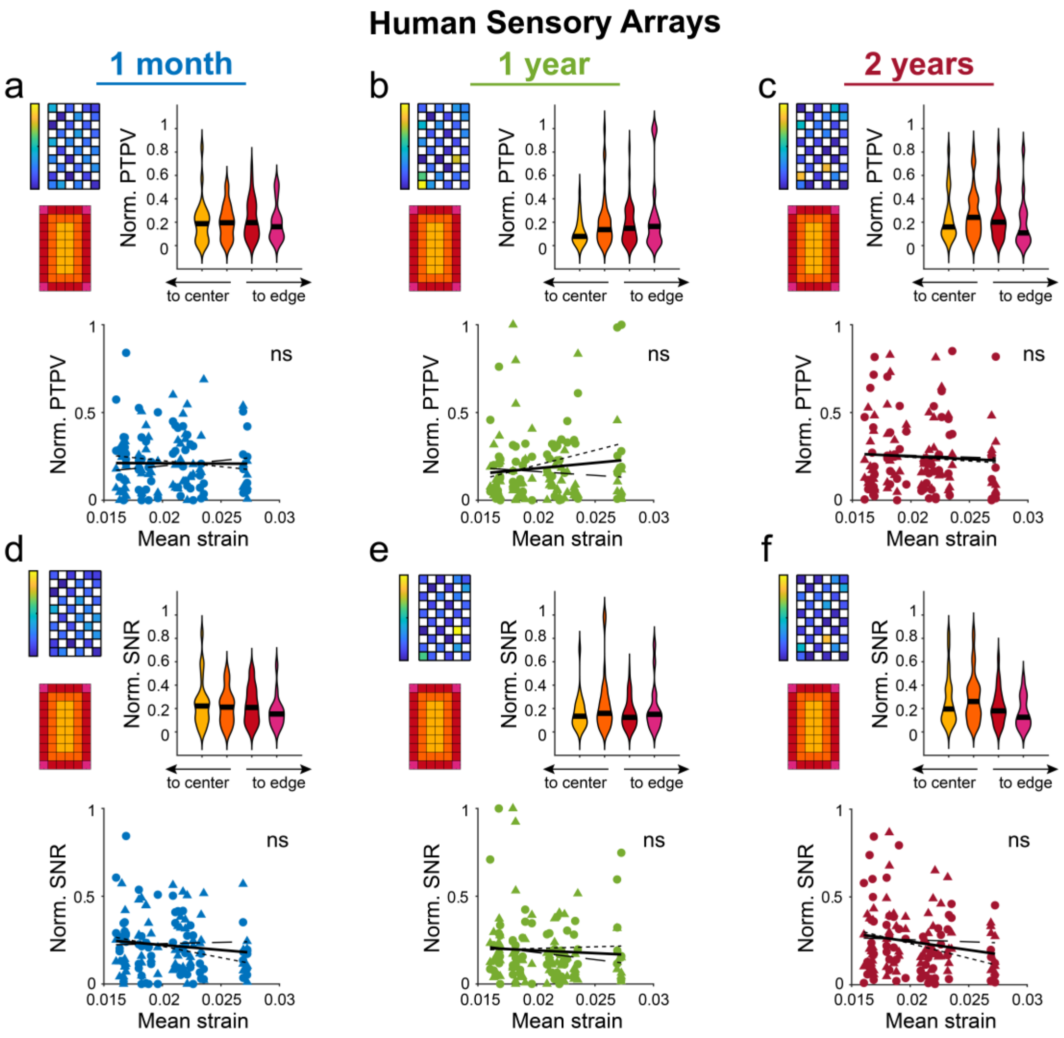


**Supplementary Figure 9. Spontaneous peak-to-peak voltage and SNR from implanted human somatosensory arrays show limited relationship with micromotion strains.** (a-c) Normalized PTPV measured at 1-mo, 1-yr, and 2-yrs post-implantation in somatosensory cortex of human study participants (N=2 participants, n=4 arrays). Heatmap shows an example array. Violin plots show no dependence on electrode location within the array (Kruskal-Wallis test with post-hoc Dunn's test). Normalized PTPVs were correlated with modeled von Mises strains using Spearman's rank-order correlation. No significant correlation as observed. (d-f) Same as above, but using normalized SNR measured at 1-mo, 1-yr, and 2-yrs post-implantation in somatosensory cortex of human study participants. No strain correlation or edge/interior dependence was observed. Black line in violin plots show median value. P2 data is plotted with circles and small-dashed trend line. P3 data is plotted with triangles and large-dashed trend line. The thick trend line fits the combined data.
